# Age-associated repression of HNF4α is a driver and potential therapeutic target in liver metabolic dysfunction

**DOI:** 10.64898/2026.11.05.764445

**Authors:** Jessica Proulx, Kathryn Lande, Xue Lei, Aaron P. Havas, Marcos Garcia Teneche, Adarsh Rajesh, Hiroshi Tanaka, Michael A LaPorta, K. Garrett Evensen, Sainath Mamde, Lu Wang, Armin Gandhi, Rouven Arnold, Andrew Davis, Angela Liou, Rabi Murad, Guillermina Garcia, Monica Sevilla, Rebecca Porritt, Angelo D’Alessandro, Shelley L. Berger, Allen Wang, Kevin Y. Yip, Susan Kaech, Xianlin Han, April E. Williams, Peter D. Adams

## Abstract

Aging is a major risk factors for metabolic dysfunction-associated steatotic liver disease (MASLD). However, the mechanisms by which aging disrupts hepatocyte transcriptional networks to drive metabolic dysfunction remain poorly understood. Hepatocyte nuclear factor 4 alpha (HNF4α) is a master regulator of hepatocyte identity and metabolic homeostasis, but its role in liver aging has not been defined. Here, we perform a comprehensive multi-omic profiling of young and aged mouse livers combined with targeted HNF4α functional studies to define how aging remodels HNF4α-dependent gene regulation and whether HNF4α can be a target to reverse age-associated liver dysfunction. Despite preserved HNF4α expression with age, aging was associated with reduced HNF4α chromatin occupancy and transcriptional activity, accompanied by epigenetic remodeling, including changes in histone acetylation, chromatin accessibility, and DNA methylation. Integration and comparison of HNF4α CUT&RUN with hepatocyte-specific *Hnf4a* deletion in young and aged livers identified high-confidence overlapping but distinct young- and old-specific HNF4α target gene networks. In young hepatocytes, HNF4α coordinates two complementary transcriptional programs: activation of genes governing hepatocyte identity, lipid metabolism, and metabolic function, while simultaneously inhibiting alternative developmental, proliferative, and inflammatory programs. Spatial transcriptomic analyses further revealed zone-specific specialization of HNF4α-dependent gene programs. Aging selectively disrupted these regulatory networks, resulting in compromised liver identity and metabolic gene programs, de-repression of inflammatory pathways and alternative cellular identity programs and disrupted hepatocyte zonation, consistent with age-dependent lipidomic and metabolic changes, hepatic fat accumulation, and immune infiltration. Hepatocyte-specific *Hnf4a* deletion in young mice recapitulated key features of liver aging, including impaired hepatocyte identity, metabolic dysfunction, steatosis, inflammatory activation and altered hepatic immune-cell composition, while deletion in aged mice further exacerbated steatosis, inflammation, and hepatic dysfunction. Conversely, hepatocyte-specific HNF4α restoration in aged mice significantly reduced hepatic steatosis and lipid accumulation, restored fatty acid metabolic programs, and partially reversed age-associated transcriptional changes. However, inflammatory and interferon-associated programs were not uniformly reversed, revealing differential responsiveness of metabolic and inflammatory pathways to HNF4α restoration. Together, these findings identify age-associated impairment of HNF4α activity as a functional contributor to liver aging, linking dysregulation of hepatocyte identity, metabolic homeostasis, and inflammatory control to age-associated hepatic dysfunction. Importantly, the selective reversibility of metabolic defects following HNF4α restoration establishes HNF4α as a promising strategy for preserving hepatocyte homeostasis and reversing age-associated hepatic metabolic dysfunction.

## Background

Aging is a major risk factor for chronic diseases including cancer, cardiovascular disease, neurodegeneration, and metabolic disorders such as type II diabetes and metabolic dysfunction-associated steatotic liver disease (MASLD)^1–3^. The liver is a central regulator of systemic metabolic homeostasis, coordinating lipid and glucose metabolism, nutrient storage, bile acid synthesis, and xenobiotic detoxification. With age, the liver undergoes progressive metabolic remodeling characterized by increased lipid accumulation, altered mitochondrial and lipid metabolism, and chronic low-grade inflammation which overlap with features of MASLD and may increase susceptibility to metabolic liver disease and its complications^3–6^. MASLD is now the most prevalent chronic liver disease worldwide, yet effective interventions for advanced MASLD remain limited^7–9^. As aging increases susceptibility and severity to progressive metabolic liver disease, defining how aging disrupts hepatocyte function may reveal therapeutically targetable vulnerabilities that precede irreversible liver disease.

Aging is increasingly recognized as a state of progressive epigenetic and chromatin remodeling, including changes in DNA methylation, histone modifications, chromatin architecture, and transcription-factor activity that can compromise the fidelity of cell-type-specific gene expression programs required for healthy cellular function and identity^10–14^. Consistent with this idea, liver aging is associated with extensive chromatin remodeling, including altered regulation of hepatocyte-specific and developmental gene programs^10, 13, 15, 16^. Conversely, longevity-promoting interventions can suppress and even reverse age-associated epigenetic changes in the liver to partially restore youthful hepatic gene networks including restoring metabolic gene regulation and suppressing inflammatory programs^10, 13, 16^. These observations suggest that age-associated liver dysfunction reflects, at least in part, a failure to maintain essential liver transcriptional programs that preserve healthy hepatocyte function and identity, and that these networks may be a therapeutically tractable target in liver aging.

Hepatocyte nuclear factor 4 alpha (HNF4α; NR2A1) is a lineage-defining nuclear receptor and master regulator of hepatocyte identity and metabolism, controlling extensive gene networks involved in lipid, glucose, amino acid, bile acid, and xenobiotic metabolism^17^. Conditional deletion of *Hnf4a* in adult hepatocytes causes widespread transcriptional dysregulation, hepatic lipid accumulation, impaired lipid homeostasis, and loss of differentiated liver function^18^. Consistent with its central role in hepatic homeostasis, HNF4α activity declines with increasing severity across chronic liver diseases and is associated with adverse clinical outcomes^19^. HNF4α dysfunction has also been implicated in MASLD/MASH, alcohol-associated liver disease, fibrosis, cirrhosis, and hepatocellular carcinoma^20, 21^. Reduced HNF4α-dependent transcription can occur despite preserved HNF4α protein expression, with epigenetic and chromatin remodeling implicated in impaired regulatory activity ^19, 21^. Conversely, enhancing HNF4α activity can restore hepatic metabolic programs and improve pathology in preclinical models of fibrosis, steatohepatitis, and tumorigenesis^22, 23^. Although HNF4α is best established as an activator of liver-specific metabolic programs, reciprocal interactions between HNF4α and inflammatory or stress-associated signaling have been described in liver injury and chronic disease^20, 21, 24, 25^, suggesting HNF4α may also function as an important constraint for inflammatory signaling. However, the extent to which HNF4α maintains metabolic and inflammatory homeostasis in the liver have not been systematically defined. Moreover, despite substantial evidence linking HNF4α loss or dysfunction to chronic liver disease, the role of HNF4α in physiological liver aging, the mechanisms by which its regulatory network may become disrupted with age, and whether HNF4α is therapeutically targetable in aged hepatocytes remains unestablished.

Here, we define how physiological aging remodels HNF4α-dependent regulatory networks in vivo. By integrating HNF4α CUT&RUN with hepatocyte-specific *Hnf4a* knockout in young and aged liver, we identify high-confidence, directionally resolved HNF4α target genes and distinguish direct HNF4α-dependent programs from secondary consequences of HNF4α perturbation. We show that aging is accompanied by coordinated loss of HNF4α-dependent metabolic and hepatocyte-identity programs and derepression of inflammatory, tissue-remodeling, and alternative cell-state programs, despite preserved HNF4α expression. HNF4α knock out recapitulated and exacerbated key features of aged liver, including impaired hepatocyte programs, metabolic dysfunction, steatosis, immune infiltration and inflammatory activation, demonstrating that loss of HNF4α accelerates liver aging. Finally, we demonstrate that HNF4α ectopic expression can reverse select features of age-associated liver dysfunction. Together, our findings establish HNF4α as a gatekeeper of hepatocyte identity, metabolic homeostasis, and inflammatory restraint whose regulatory activity is compromised during aging. Moreover, restoration of HNF4α-dependent transcriptional programs is a promising strategy to reverse aspects of age-associated liver dysfunction.

## Results

### Age-associated liver lipidomic and metabolic dysfunction is associated with epigenetic remodeling and repression of HNF4α transcriptome

To gain insight into potential mechanisms driving age-related hepatocyte dysfunction and liver disease development, we performed comprehensive transcriptomic, lipidomic, metabolomic, and epigenomic analyses of young (4-5 months) and aged (22-23 months) male mouse livers. Our recently published RNA-seq analysis of isolated hepatocytes^26^ identified robust activation of interferon-, oncogene- and tumor suppressor-related programs in aged hepatocytes. Here, we focus on genes downregulated with age, which were enriched for core hepatocyte metabolic functions, including fatty acid (FA), bile acid, and xenobiotic metabolism, oxidative phosphorylation, and peroxisomal processes (Fig. 1A-B; FDR < 0.05; Log2FC < -0.32). Lipidomic profiling identified significant age-associated remodeling across glycerolipid, phospholipid (PL), FA, and steroid metabolic and biosynthetic pathways (Fig. 1C-D; Supp. Fig. 1A-B). Briefly, increases in triacyclglycerols (TAGs) and cholesteryl esters (CEs) were consistent with hepatic steatosis while accumulation of free FAs, FA esters of hydroxy FAs (FAHFAs), and acylcarnitine FAs (mitochondrial-associated FAs primed for oxidation) were consistent with impaired FA metabolism. Integration of hepatocyte transcriptomic data with our previously published metabolomic data^26^ further demonstrated coordinated disruption of core hepatocyte metabolic functions, including lipid and membrane metabolism, central carbon metabolism, redox and detoxification capacity, and amino-acid biosynthesis (Supp. Fig. 1C).

**Figure 1.**
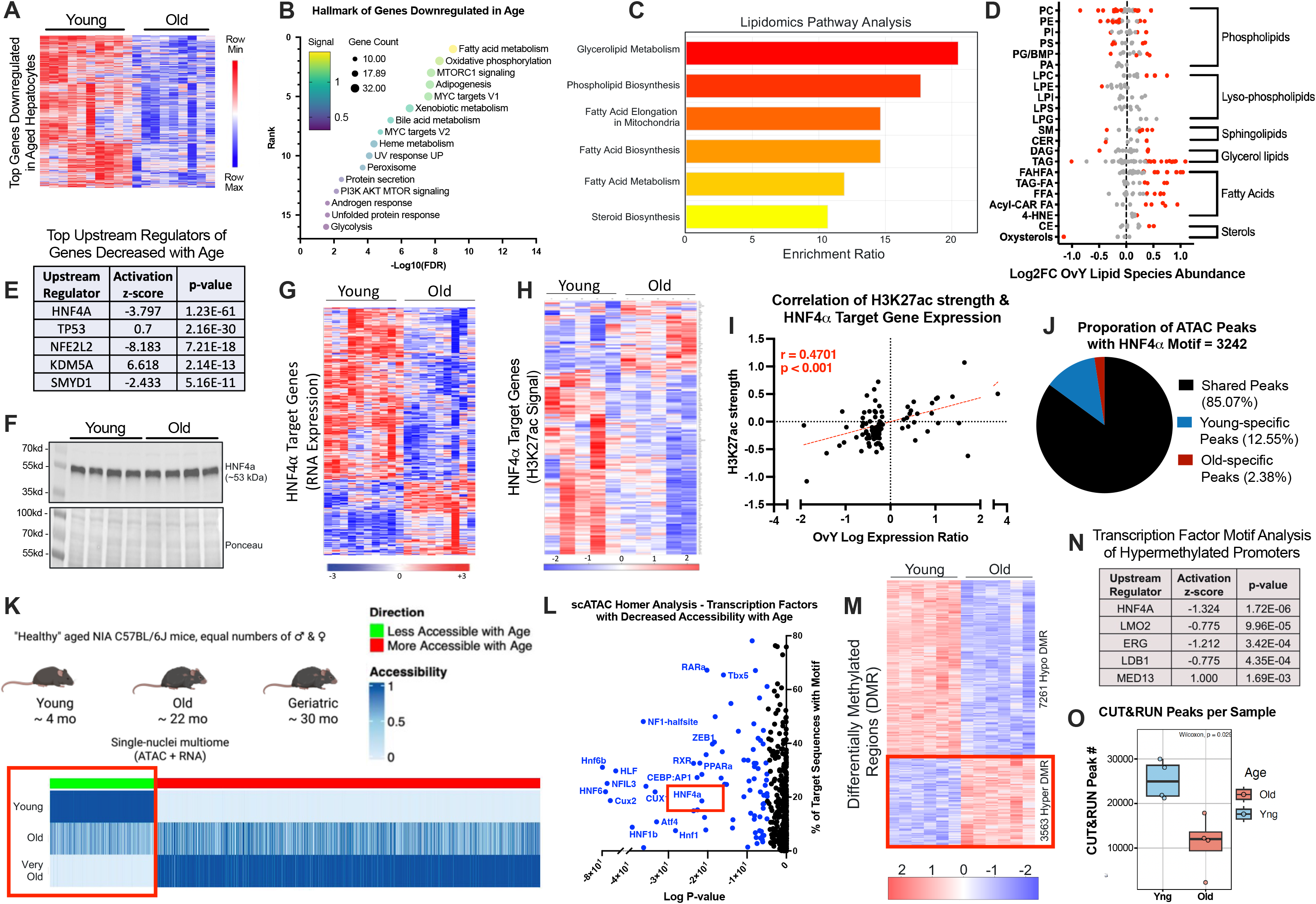
Age-associated liver lipidomic and metabolic dysfunction is associated with epigenetic remodeling and repression of HNF4α transcriptome. **A**) Heatmap of top 999 genes downregulated in old (O) versus young (Y) hepatocytes by RNA-seq (|Log2FC| > 0.32; FDR < 0.05). **B**) Hallmark pathway enrichment analysis of age-downregulated genes in A. **C**) Lipidomic pathway enrichment analysis of differentially abundant lipid species in O versus Y whole livers (p < 0.05). **D**) Log2FC between O versus Y of individual lipid species grouped by lipid class. Red dots indicate significantly altered lipid species (p < 0.05). **E**) Ingenuity Pathway Analysis (IPA) of upstream regulators associated with age-downregulated genes in A. F) Whole liver western blot (WB) of HNF4α. **G**) Heatmap of 510 IPA-defined HNF4α target genes differentially expressed (DE) between Y and O hepatocytes. **H**) Heatmap of H3K27ac signal intensity at IPA-defined HNF4α target genes shown in G with detectable H3K27ac peaks (n = 100 genes). **I**) Pearson correlation between age-associated changes in gene expression and H3K27ac signal intensity for genes included in both G and H. **J**) Proportion of HNF4α motif-containing ATAC-seq peaks classified as shared, young-specific, or aged-specific in bulk ATAC-seq of young and aged whole livers (n = 3,242 peaks) **K**) Schematic for snATAC-seq analyses in Y, O, and very O male & female hepatocytes, with heatmap illustrating progressive age-associated changes in chromatin accessibility. **L**) HOMER transcription factor motif enrichment analysis of regions exhibiting progressively decreased chromatin accessibility with age, as highlighted with red box in K. **M**) Heatmap of differentially methylated regions (DMR) in aged hepatocytes from whole genome bisulfite sequencing (WGBS) analysis (7261 hypo-DMR; 3563 hyper-DMR). **N**) Transcription factor motif analysis of hypermethylated promoters identified from M. **O**) Quantification of HNF4α CUT&RUN peak numbers in young and aged mouse livers (n = 4 per age group).

To identify transcriptional regulators associated with the age-dependent loss of hepatocyte metabolic function, we performed Ingenuity Pathway Analysis (IPA) of downregulated genes. HNF4α was the top predicted upstream regulator inhibited in aged hepatocytes (Fig. 1E; z-score = -3.797; p = 1.23E-61), These IPA-identified HNF4α-associated genes among the most highly expressed genes in hepatocytes and enriched for amino-acid, lipid, and protein metabolism (Supp. Fig. 1D-E), consistent with the established role of HNF4α in maintaining hepatocyte identity and metabolic function^17, 18^. Despite the predicted inhibition of HNF4α activity, HNF4α RNA and protein expression were unchanged between young and aged livers (Fig. 1F; Supp. Fig. 1F), and analysis of *Hnf4a* transcript usage revealed no age-dependent isoform switching (data not shown). To interrogate the broader age-dependent transcriptional response regulated by HNF4α, we repeated IPA using all genes differentially expressed (DE) with age, resulting in 510 IPA-identified HNF4α-regulated genes DE with age, including 328 downregulated and 182 upregulated genes (Fig. 1G).

We next examined whether changes in HNF4α-associated genes were accompanied by the age-associated remodeling of the regulatory chromatin landscape. Among IPA-defined HNF4α target genes with detectable H3K27ac peaks, age-associated changes in gene expression significantly correlated with changes in H3K27ac signal intensity at corresponding genomic regions (Fig. 1G-I). Consistent with these findings, H3K27ac HiChIP also supported reduced promoter-enhancer interactions at HNF4α-regulated genes with age (Supp Fig. 1G). Bulk ATAC-seq analysis showed that 85% of accessible HNF4α motif-containing regions were shared between young and aged livers; however, approximately 13% were unique to young livers and significantly enriched compared to both random (p = 8.46e-3) and aged-specific sequences (p = 1.093e-3, Fig. 1J). Consistent with this finding, single-nuclei chromatin accessibility analysis identified HNF4α-associated regulatory elements among the most significantly less accessible regions in hepatocytes undergoing progressive age-associated chromatin silencing in both male and female mice (Fig. 1K-L). Lastly, despite an overall loss of DNA methylation with age, age-associated hypermethylated promoters were selectively enriched for HNF4α motifs, with HNF4α identified as the top transcription factor associated with these regions (Fig. 1M-N). To confirm whether these chromatin regulatory changes were accompanied by altered HNF4α chromatin binding, we performed HNF4α CUT&RUN in young and aged livers. Indeed, global HNF4α chromatin occupancy was reduced in aged liver (Fig. 1O), with decreased binding observed across promoters, gene bodies, and distal regulatory regions (Supp Fig. 1H).

Altogether, these analyses identified broad age-associated disruption of hepatocyte metabolic programs accompanied by lipidomic changes consistent with steatosis and altered FA metabolism. HNF4α emerged as the top predicted upstream regulator of these transcriptional changes, with reduced chromatin occupancy and transcriptional output despite preserved HNF4α expression. Concordant changes in H3K27ac abundance, chromatin accessibility, and DNA methylation link age-associated epigenetic remodeling to impaired HNF4α-dependent transcription and loss of hepatocyte metabolic and lipid homeostasis.

### Integrated HNF4α chromatin occupancy and hepatocyte-specific genetic perturbation define high-confidence HNF4α target gene networks in young and aged liver

The HNF4α-associated genes identified in Figure 1 were primarily inferred from motif-based predictions and the curated IPA database, without considering age as a variable. We therefore sought to experimentally define high-confidence, directionally resolved HNF4α-dependent targets based on direct HNF4α binding and functional dependency in young and aged liver. To assess functional dependency, we generated a hepatocyte-specific *Hnf4a* knockout (KO) using AAV8-TBG-saCas9/U6-sgHNF4α and confirmed efficient HNF4α depletion at the DNA and protein levels three weeks after viral delivery (Fig. 2A-B; Supp. Fig. 2A). RNA-seq analysis of *Hnf4a* KO versus age-matched *Rosa26* KO control livers integrated with HNF4α CUT&RUN of young and old mouse liver identified HNF4α target genes, defined as DE following *Hnf4a* deletion and associated with HNF4α occupancy within the gene body, promoter, and/or an intergenic region within 5 kb of the transcriptional start site. Target genes were independently defined in both young and old livers (Fig. 2C). This approach identified 3,394 high-confidence HNF4α target genes in young liver and 1,155 in aged liver, with 1,032 shared between young and aged liver, 2,362 unique to young, and 123 unique to aged liver (Fig. 2D). Target genes that decreased following *Hnf4a* deletion were classified as HNF4α-activated targets, whereas genes that increased were classified as HNF4α-inhibited targets. Averaged CUT&RUN coverage profiles demonstrate chromatin occupancy at representative HNF4α-activated and -inhibited target genes, with corresponding gene expression changes following Hnf4a deletion (Fig. 2E-F).

**Figure 2.**
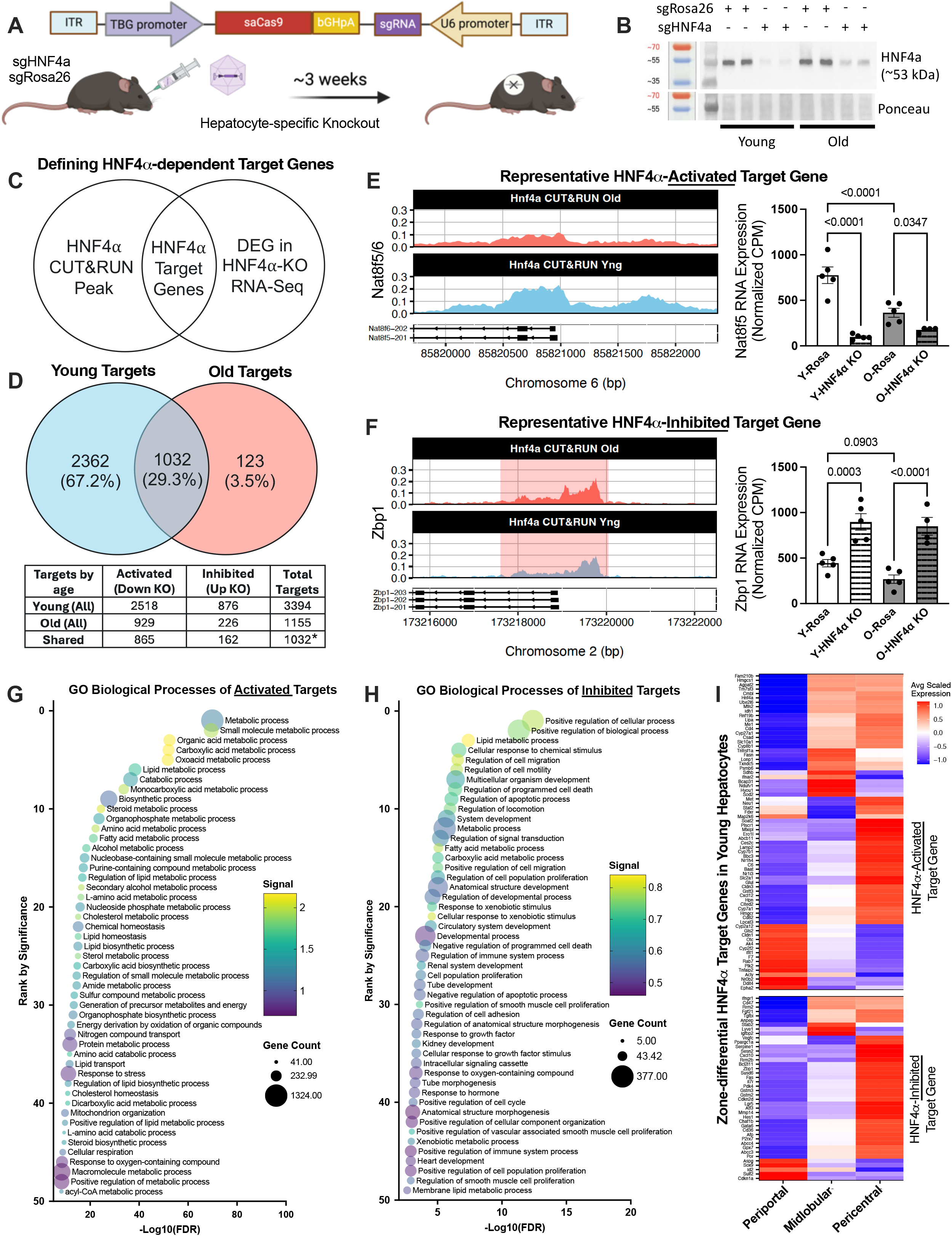
Integrated HNF4α chromatin occupancy and hepatocyte-specific genetic perturbation define high-confidence HNF4α target gene networks in young and aged liver. **A**) Schematic of hepatocyte-specific *Hnf4a* knockout (KO) using AAV-mediated delivery of sgRNAs targeting Hnf4a or Rosa26 (control), with liver collection approximately 3 weeks after administration. **B**) WB confirming reduced HNF4α protein expression following *Hnf4a* KO in Y and O mice. **C**) Schematic illustrating the identification of HNF4α target gene networks through integration of HNF4α CUT&RUN chromatin occupancy and differential gene expression following hepatocyte-specific Hnf4a deletion. **D**) Venn diagram illustrating the overlap of Y- and O-HNF4α target genes, with corresponding numbers for HNF4α-activated (downregulated following KO) and HNF4α-inhibited (upregulated following KO) target genes (*Five shared target genes exhibited opposing regulatory classifications between Y and O). **E-F**) Averaged, bins-per-million (BPM)-normalized HNF4α CUT&RUN coverage tracks across biological replicates within each age group, with corresponding hepatocyte RNA-seq expression following *Hnf4a* KO, for representative **E**) HNF4α-activated target gene (*Nat8f5*) and **F**) HNF4α-inhibited target gene (*Zbp1*). Gene Ontology Biological Process (GO:BP) enrichment analyses of **G**) Y-HNF4α-activated target genes and **H**) Y-HNF4α-Inhibited target genes. **I**) Heatmap of zone-differential Y-HNF4α target gene expression in WT hepatocytes using MERFISH pseudobulk analysis (n = 2 young livers), stratified by Y-HNF4α-activated and Y-HNF4α-inhibited target genes.

We next characterized the biological functions of HNF4α target genes in young adult liver. HNF4α-activated targets were broadly enriched for core metabolic and biosynthetic processes, including small molecule, organic and carboxylic acid, lipid and amino-acid processes (Fig. 2G; Supp Fig 2B). While HNF4α-inhibited targets were also enriched for metabolic processes, including lipid metabolism, they additionally were enriched for alternative developmental, proliferative, migratory, and apoptotic processes as well as immune system processes (Fig. 2H; Supp. Fig. 2C). Together, these results support a dual regulatory role for HNF4α in simultaneously promoting and suppressing distinct metabolic programs while restraining alternative cellular programs.

Because hepatocyte identity and metabolic function are spatially organized according to distinct liver zones, we next examined the distribution of HNF4α target genes at single-cell resolution using a custom 500-gene MERFISH spatial transcriptomics panel. Of the 126 HNF4α-dependent target genes represented in our panel (81 HNF4α-activated and 45 HNF4α-inhibited targets), 114 were preferentially enriched in at least one hepatocyte zone in young liver with distinct HNF4α target signatures across periportal, midlobular, and pericentral hepatocytes (Fig. 2I). Notably, the most pronounced differences observed were between periportal and pericentral populations, including canonical zonation-associated markers Cyp2f2 in periportal hepatocytes and Glul in pericentral hepatocytes, consistent with zonal HNF4α specialization in young livers.

Collectively, integration of HNF4α occupancy with hepatocyte-specific *Hnf4a* KO defined high-confidence, directionally resolved HNF4α target gene networks in young and aged hepatocytes. In young, this network encompasses complementary programs that promote differentiated hepatocyte metabolic function and identity while simultaneously suppressing alternative cellular and developmental programs, including proliferation, cell stress, inflammation, and aberrant signaling and metabolic processes. Moreover, HNF4α-dependent programs in young healthy liver display substantial spatial organization patterns across hepatocyte zones, supporting a role for HNF4α in contributing to the functional specialization of distinct hepatocyte populations.

### Age-associated remodeling of HNF4α target genes disrupts hepatocyte identity and metabolic programs and permits inflammation

Having analyzed HNF4α activated and inhibited target genes in young adult liver, we next integrated high-confidence HNF4α-dependent regulatory programs with young-versus-aged hepatocyte RNA-seq from Figure 1 to determine how these programs change with age and their associated biological functions. HNF4α-activated target genes, defined in both young and aged liver, were significantly enriched among genes downregulated with age and depleted among genes upregulated with age. Conversely, HNF4α-inhibited targets defined in young liver were enriched for genes upregulated with age (Fig. 3A).

**Figure 3.**
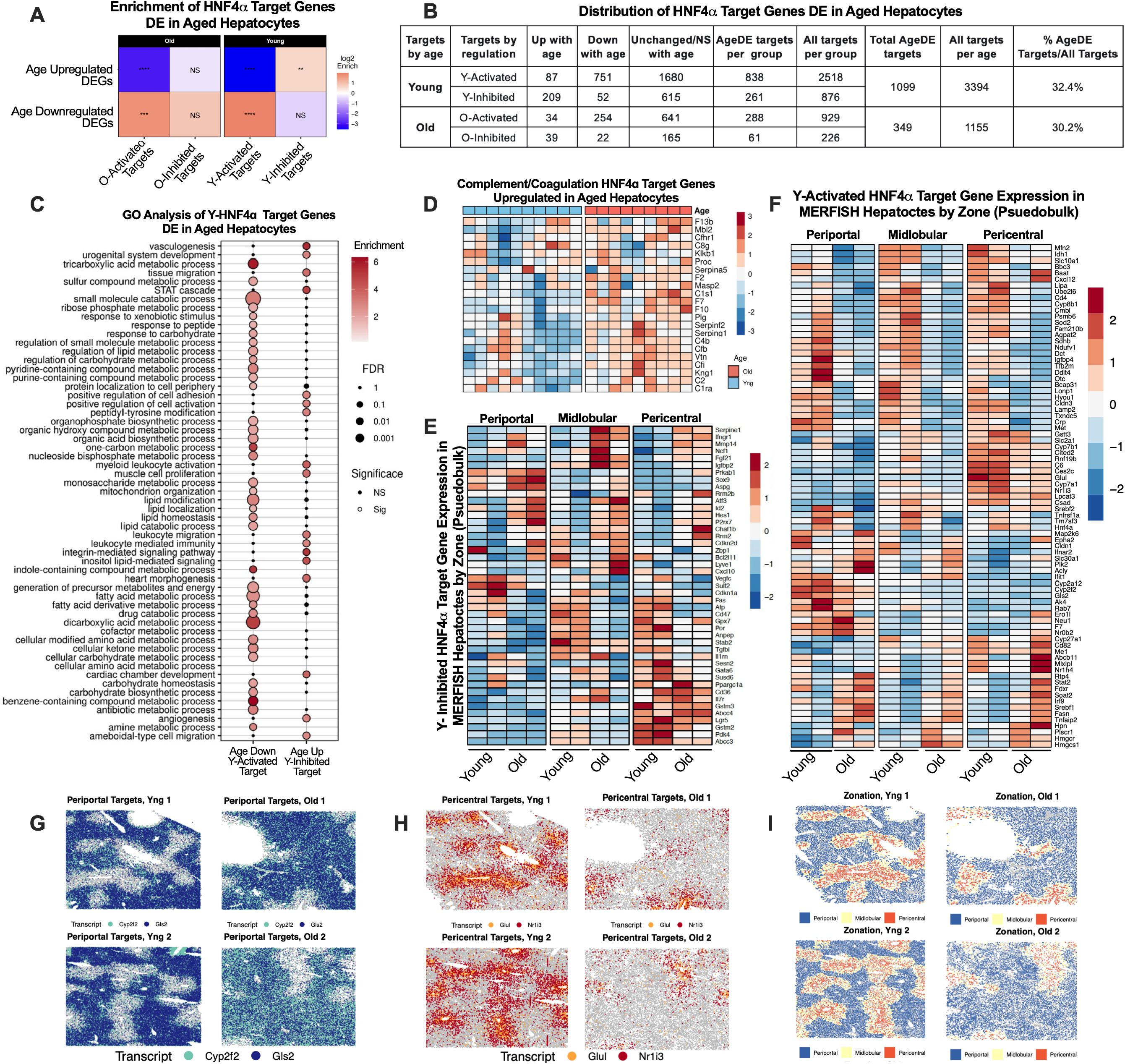
Age-associated remodeling of HNF4α target genes disrupts hepatocyte identity and metabolic programs and permits inflammation. **A**) Enrichment scores for Y- and O-HNF4α target genes DE in WT aged hepatocyte RNA-seq. **B**) Distribution of Y- and O-HNF4α target genes according to their regulatory classification and age-associated differential expression in WT aged hepatocyte RNA-seq, including the number of upregulated, downregulated, and unchanged target genes. **C**) GO:BP enrichment analysis of Y-HNF4α-activated targets downregulated with age and Y-HNF4α-inhibited targets upregulated with age. **D**) Heatmap of complement- and coagulation-associated Y-HNF4α-activated target genes upregulated with age in hepatocyte RNA-seq. **E-F**) Heatmaps of pseudobulk MERFISH gene expression in Y and O WT hepatocyte for all detected **E**) Y-inhibited and **F**) Y-activated HNF4α target genes, stratified by periportal, midlobular, and pericentral zones (nYng=2; nOld=2). **G-H**) Representative MERFISH spatial expression maps illustrating age-associated changes in selected zone-enriched HNF4α target genes, including **G**) periportal-associated *Cyp2f2* and *Gls2* and **H**) pericentral-associated *Glul* and *Nr1i3*. Images are shown for two biological replicates per age group. **I)** Corresponding MERFISH spatial maps illustrating hepatocyte zonation by periportal, midlobular, and pericentral classifications in Y and O livers.

To better dissect the extent, directionality, and selectivity of age-associated remodeling across the HNF4α target network, we categorized HNF4α-dependent programs according to age-differential expression (i.e. upregulated, downregulated, or unchanged/NS; Fig 3B). Among the 3,394 HNF4α targets defined in young liver, 1,099 (32.4%) were DE in aged hepatocytes, and of these 1,099 genes the vast majority (∼87%) were either age-downregulated HNF4α-activated targets (751 genes) or age-upregulated HNF4α-inhibited targets (209 genes). Young HNF4α-activated targets that decreased with age were enriched for core hepatocyte and metabolic processes, including FA and lipid metabolism, lipid catabolism/modification, carbohydrate and amino acid metabolism, mitochondrial function, nucleotide metabolism, and xenobiotic/drug metabolism (Fig. 3C). In contrast, young HNF4α-inhibited targets that increased with age were enriched for angiogenesis/vasculogenesis, leukocyte and myeloid activation, cell adhesion and integrin signaling, proliferation, migration, STAT signaling, and alternative developmental and morphogenetic programs (Fig. 3C).

Notably, the majority (67%) of young HNF4α-activated targets were unchanged/NS with age (1680 out of 2518 genes) and encompassed a wide range of metabolic and hepatocyte functions, indicating that while many HNF4α-dependent metabolic programs are disrupted with age, a substantial fraction remains relatively stable (Supp. Fig. 3A). In contrast, a small subset of young HNF4α-activated targets (87 genes) increased with age and were enriched for protein activation cascade pathways, specifically complement- and coagulation-associated genes (Fig. 3D; Supp. Fig. 3A). Interestingly, DE young HNF4α-inhibited targets (261 genes) were enriched for multiple biological processes represented across both age-upregulated and age-downregulated categories; however, select processes including angiogenesis/vasculogenesis, leukocyte and myeloid activation, integrin signaling, and STAT cascade were exclusively represented in the age-upregulated targets (209 genes; Supp. Fig. 3B). Altogether, HNF4α target genes upregulated with age (both activated and inhibited) are enriched for immune related terms (complement, leukocyte and myeloid activation, STAT cascade genes).

We next examined single-cell age-associated remodeling of HNF4α target genes across hepatocyte zones using spatial MERFISH. Of the 126 HNF4α target genes represented in the MERFISH panel, 122 exhibited significant age-associated changes in at least one hepatocyte zone (Fig. 3E-F; Supp. Fig. 3C-F). Notably, while age-associated changes were uniform across hepatocyte zones, a subset of genes exhibited zone-specific transcriptional responses (Supp. Fig. 3C-F). Consistent with spatial remodeling, canonical zonation-associated HNF4α targets exhibited altered expression patterns with age (Fig. 3G-I). Periportal targets Cyp2f2 and Gls2 exhibited an expanded or relatively increased spatial expression pattern with age (Fig. 3G), whereas pericentral HNF4α targets Glul and Nr1i3 showed a reduced expression pattern with age (Fig. 3H). These findings suggest that age-associated remodeling of HNF4α-dependent transcription may be a key feature of the recently but robustly reported age-associated disruption of normal hepatocyte zonation (Fig. 3I)^26–28^.

Collectively, aging substantially and selectively remodels HNF4α-dependent transcriptional programs. The predominant changes include loss of HNF4α-activated metabolic and hepatocyte identity programs together with a derepression of alternative identity and inflammatory cellular programs. These changes extend across distinct hepatocyte populations and are spatially heterogeneous, including alterations in zone-associated HNF4α programs and disruption of canonical hepatocyte zonation.

### Hepatocyte-specific loss of HNF4α recapitulates and exacerbates key features of liver aging

Given the predominant age-associated decline in HNF4α activity, we next investigated whether loss of HNF4α is sufficient to recapitulate or exacerbate key features of liver aging. To address this, we used AAV8-TBG-saCas9/U6-sgHNF4α for hepatocyte-specific *Hnf4a* deletion in young and aged mice and analyzed the resulting transcriptional, histological, and immune phenotypes. Loss of HNF4α induced extensive transcriptional remodeling in young and aged mice, with 8,739 genes DE following *Hnf4a* deletion in young mice and 5,857 in aged mice (Fig. 4A). Principal component analysis demonstrated clear separation by both HNF4α KO and age (Fig. 4B). To determine the extent to which HNF4α loss recapitulates age-associated transcriptional changes, we compared the transcriptional response to *Hnf4a* deletion in young liver with that observed during physiological aging in wild-type (WT) hepatocytes. Transcriptional changes induced by *Hnf4a* deletion significantly correlated with age-associated transcriptional changes in WT hepatocytes, with 63.3% of age-upregulated genes (1,467/2,319) and 62.7% of age-downregulated genes (1,194/1,904) similarly DE following *Hnf4a* deletion (Fig. 4C; Supp. Fig. 4A–B).

**Figure 4.**
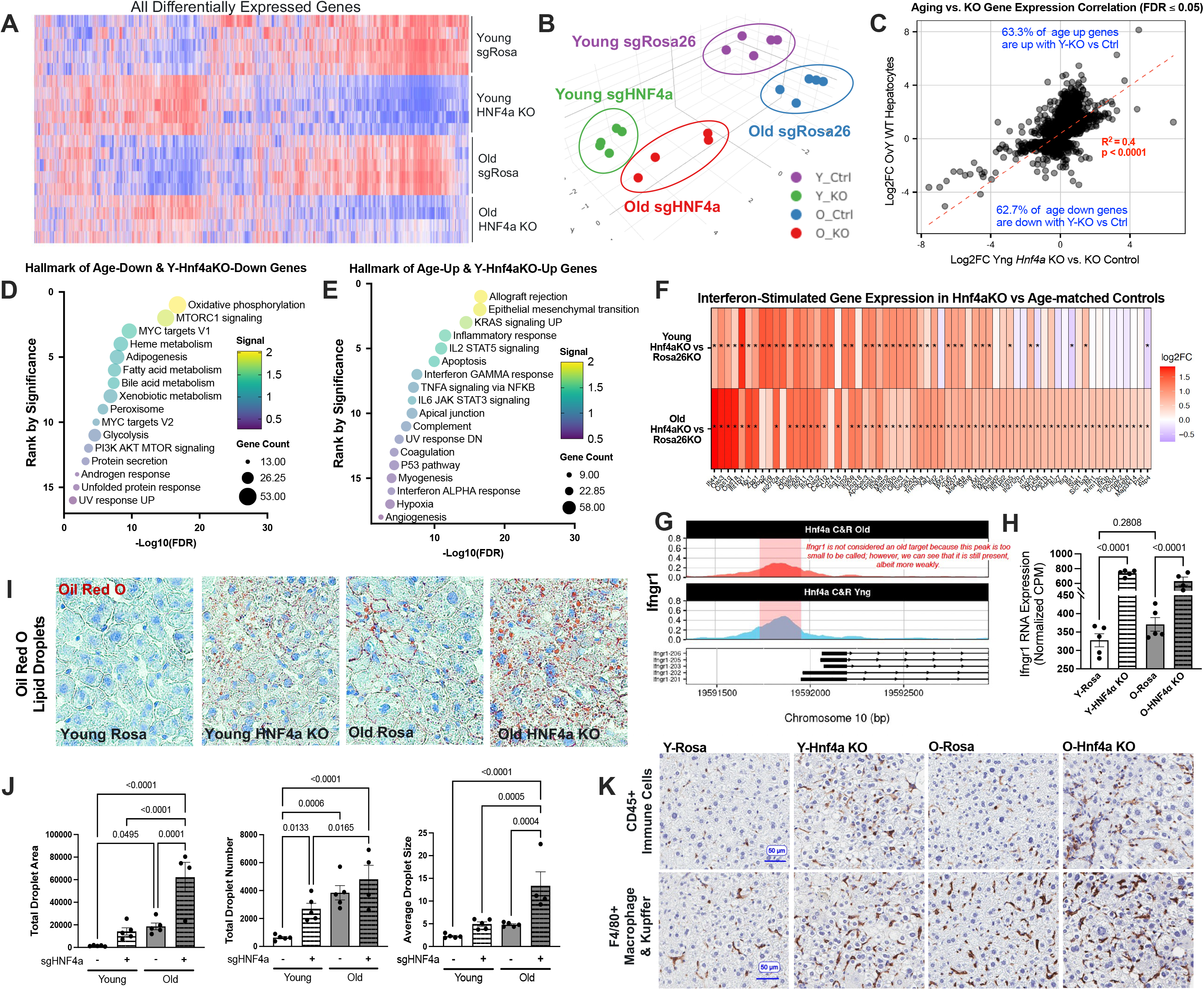
Hepatocyte-specific loss of HNF4α recapitulates and exacerbates key features of liver aging. **A**) Heatmap of all liver DE genes identified by whole liver RNA-seq following hepatocyte-specific *Hnf4a* KO in Y and O mice. **B**) Three-dimensional principal component analysis plot for each mouse in RNA-seq. **C**) Correlation of log2FC gene expression changes between physiological aging (O vs Y WT hepatocytes; FDR ≤ 0.05) and hepatocyte-specific *Hnf4a* deletion (Y-KO vs Y sgRosa26 controls; FDR ≤ 0.05). Pearson correlation and linear regression analyses were performed. **D-E**) Hallmark pathway enrichment analyses of genes concordantly regulated by physiological aging and Y hepatocyte-specific *Hnf4a* deletion, including **D**) genes downregulated in both comparisons (lower left quadrant in C; 1,194 genes) and **E**) genes upregulated in both comparisons (upper right quadrant in C; 1,467 genes). **F**) Heatmap of log2FC gene expression of interferon-stimulated genes (ISGs) significantly upregulated following *Hnf4a* KO in Y or O versus age-matched controls. **G**) Averaged, bins-per-million (BPM)-normalized HNF4α CUT&RUN coverage track and **H**) KO RNA-seq gene expression plot for *Ifngr1*. **I**) Representative Oil Red O (ORO)-stained liver images illustrating hepatic lipid accumulation following *Hnf4a* deletion in Y and O mice. **J**) Quantification of ORO-positive lipid droplets, including total droplet area, droplet number, and average droplet size. **K**) Representative liver immunohistochemistry images for CD45+ (immune cells) and F4/80+ (macrophages & Kupffer cells) in Y and O *Hnf4a* KO and age-matched control mice.

We next characterized the age-associated biological programs altered by HNF4α loss, which encompasses both direct HNF4α targets from Figure 3 and the secondary/indirect programs induced by *Hnf4a* KO. Concordantly downregulated genes were enriched for core hepatic metabolic processes, including pathways identified in our age-associated transcriptomic and metabolomic analyses (Fig. 4D; Supp. Fig. 4C). Conversely, genes concordantly induced following Hnf4a deletion were enriched for inflammatory and immune-associated pathways (Fig. 4E Supp. Fig. 4D). Consistent with activated interferon signaling during liver aging^26^ *Hnf4a* deletion robustly induced interferon-stimulated genes (ISGs) and interferon-associated pathways in both young and aged livers (Fig. 4E-F; Supp Fig 4E). Integration with HNF4α CUT&RUN identified direct occupancy at a subset of ISGs, including a major upstream regulator of interferon signaling, Ifngr1, which is a direct HNF4α-inhibited target in *young* livers (Fig. 4G-H; Supp. Fig. 4F).

We performed Oil Red O staining (ORO) to assess hepatic lipid droplet accumulation and immunohistochemistry for select immune cell markers. ORO staining revealed increased hepatic lipid droplet number, mean droplet size, and total droplet area following *Hnf4a* deletion in young mice, with exacerbated lipid droplet accumulation and steatosis in aged KO mice livers (Fig. 4I-J). Immunohistochemistry analyses demonstrated increased CD45⁺ immune cells, F4/80⁺ macrophages/Kupffer cells and B220⁺ B cells following *Hnf4a* deletion. Interestingly, there was also a trend towards decreased CD3⁺ T cells (Fig 4K; Supp Fig. G-H).

To further characterize changes in hepatic immune-cell composition, we performed flow cytometry profiling (Supp. Fig. 4I-N). *Hnf4a* deletion showed trends towards reduced total CD3⁺ T cells frequencies, including CD8⁺ T cells among CD45^hi^ immune cells, in both young and aged livers, despite increased trends between normal aging (Supp. Fig. 4I, L). In young livers, *Hnf4a* deletion significantly decreased conventional CD4⁺ T cell frequencies (CD4⁺ FoxP3^-^) while increasing CD4⁺ regulatory T cells (Tregs) and B cells frequencies (Supp. Fig. 4J, K, N). These changes in conventional CD4⁺ T cell and Treg frequencies paralleled those observed during physiological aging. Conversely, HNF4α loss increased macrophage frequencies in aged KO livers relative to their age-matched controls, with a non-significant trend in young KO livers (Supp. Fig. 4M); a change not observed during physiological aging.

Collectively, these findings demonstrate that hepatocyte HNF4α is required to maintain hepatic metabolic programs while restraining inflammatory and immune-associated programs. Loss of HNF4α in young liver recapitulated substantial molecular and physiological features of aging, including hepatocyte metabolic gene repression, hepatic steatosis, inflammatory activation, and some features of the aged hepatic immune environment. Importantly, *Hnf4a* deletion in aged liver exacerbated hepatic dysfunction, demonstrating that HNF4α remains functionally important despite its age-associated suppression. Together, these findings identify impaired HNF4α activity as a functional contributor to age-associated liver dysfunction while highlighting the continued importance of residual HNF4α activity in preserving hepatocyte identity and metabolic homeostasis during aging.

### Hepatocyte-specific restoration of HNF4α reverses age-associated liver steatosis and restores FA metabolism gene expression

Our findings demonstrate that HNF4α transcriptional activity is reduced during liver aging and that loss of HNF4α is sufficient to recapitulate key features of the aged liver. We therefore asked whether restoring HNF4α activity in aged hepatocytes could reverse age-associated hepatic dysfunction. To test this, we ectopically expressed HNF4α in hepatocytes of young and aged mice using AAV8-TBG-*Hnf4a*, with AAV8-TBG-mCherry as control. Mice were culled and tissues collected and examined for hepatic histology, lipid composition and gene expression, three weeks after virus infection (Fig. 5A). Successful ectopic expression was validated by RNA and protein expression (Supp. Fig. 5A-C)

**Figure 5.**
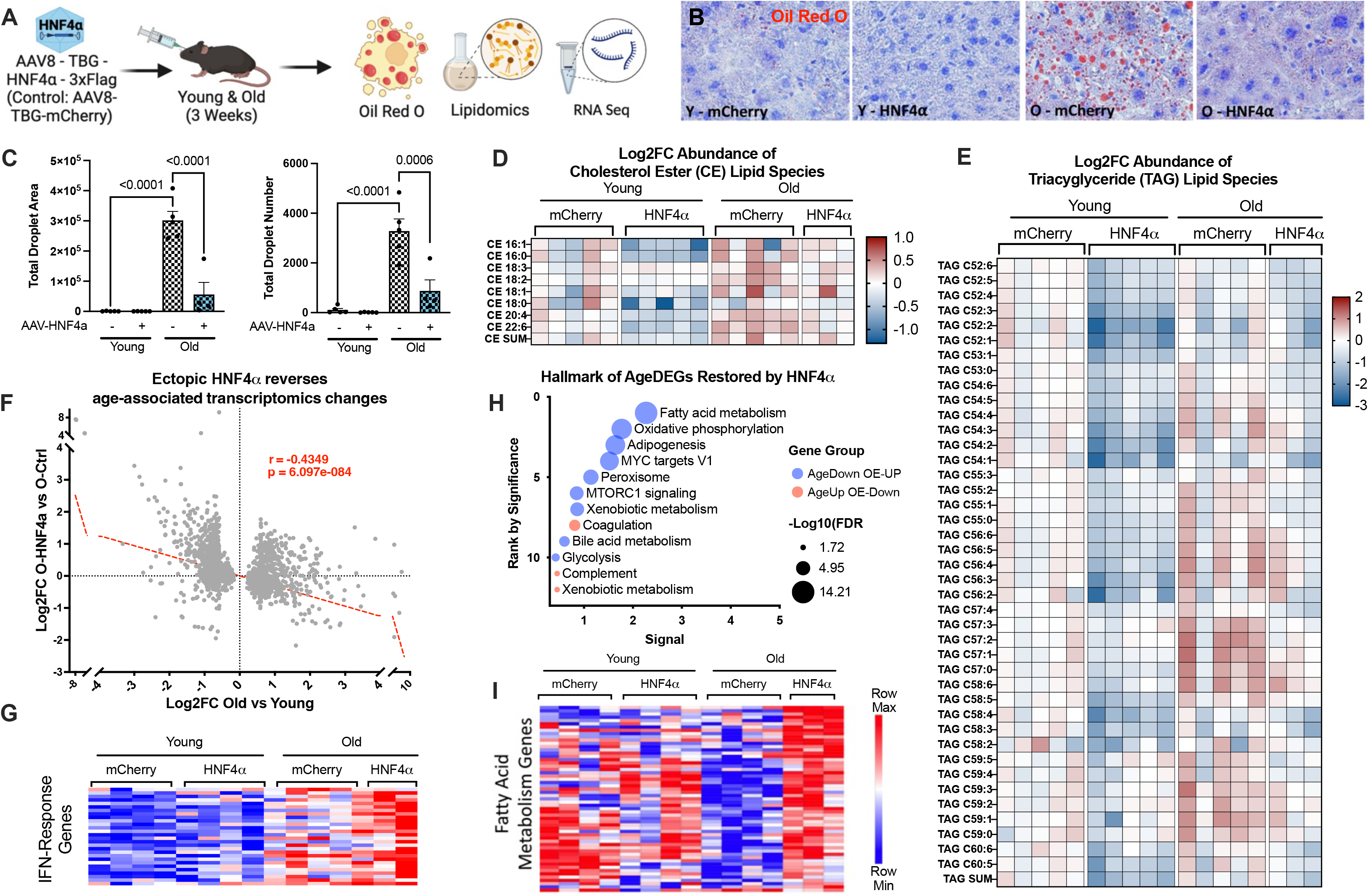
Hepatocyte-specific restoration of HNF4α reverses age-associated liver steatosis and restores FA metabolism gene expression. **A**) Schematic of hepatocyte-specific HNF4α overexpression using AAV8-TBG-HNF4α-3xFlag or AAV8-TBG-mCherry (control) in Y and O mice, with liver collection 3 weeks after administration for histological, lipidomic, and transcriptomic analyses. **B**) Representative ORO images illustrating hepatic lipid accumulation in Y and O mouse livers following AAV-mediated HNF4α overexpression or mCherry control. **C**) Quantification of ORO-positive lipid droplets, including total droplet area and droplet number. **D-E**) Heatmaps of log2FC in lipid species abundance relative to Y mCherry controls, showing **D**) cholesterol ester (CE) and **E**) triacylglycerol (TAG) species across Yand O mice treated with AAV-HNF4α or AAV-mCherry **F**) Pearson correlation plot of log2FC in gene expression between physiological aging (O vs Y control livers) and HNF4α overexpression in aged mice (O-HNF4α vs O-mCherry), based on whole-liver RNA-seq (r = -0.4349, p = 6.097 x 10^-84^). **G**) Heatmap of IFN response genes increased in O vs Y and O-HNF4α vs O-mCherry. **H**) Hallmark pathway enrichment analysis of age-associated DE genes reversed by O-HNF4α vs O-mCherry **I**) Heatmap of fatty acid metabolism-associated genes downregulated with age and increased following HNF4α overexpression in aged livers.

Consistent with findings above (Fig. 4I-J), aged control mice exhibited increased hepatic lipid accumulation compared with young controls. Strikingly, ectopic HNF4α expression significantly reversed age-associated steatosis, including lipid droplet number and area (Fig. 5B-C). We next performed untargeted lipidomic profiling to determine whether this histological improvement was accompanied by broader changes in hepatic lipid composition. HNF4α restoration significantly remodeled the liver lipidome, with global reductions in multiple lipid species associated with hepatic steatosis, including triacylglycerols (TAGs), cholesteryl esters (CEs), and total cholesterol (Fig. D-E; Supp. Fig. 5D-F). Bulk RNA-seq following hepatocyte-specific ectopic HNF4α revealed a significant inverse relationship between HNF4α restoration and age-associated transcriptional changes, indicating partial reversal of the aged liver transcriptome (Fig. 5F). Interestingly, not all age-associated gene expression changes were reversed by ectopic HNF4α. For example, inflammatory and interferon-associated programs were not uniformly suppressed. Age-upregulated interferon-response genes remained elevated in HNF4α-expressing aged livers, with selected interferon pathway components even further increased following HNF4α restoration (Fig. 5G). In contrast, age-downregulated genes restored by HNF4α re-expression were enriched for FA metabolism, mitochondrial and peroxisomal processes, and adipogenesis, whereas age-upregulated genes reversed by HNF4α restoration were enriched for coagulation, complement, and xenobiotic metabolism (Fig. 5H). Consistent with these pathway-level changes, FA metabolism genes significantly reduced with age were preferentially re-induced following HNF4α restoration (Fig. 5I).

Collectively, these findings demonstrate that restoring HNF4α activity in aged hepatocytes reverses key features of age-associated hepatic metabolic dysfunction. While age-associated inflammatory and interferon programs remained comparatively resistant to restoration, ectopic HNF4α reduced hepatic lipid accumulation, remodeled the hepatic lipidome, and restored transcriptional programs involved in FA metabolism and mitochondrial function.

## Discussion

Aging is accompanied by progressive erosion of tissue identity and metabolic homeostasis, yet the mechanisms linking age-associated chromatin remodeling to loss of differentiated cellular function remain incompletely understood. Here, we identify HNF4α as a central regulator of age-associated hepatocyte dysfunction and demonstrate that its transcriptional activity is substantially remodeled during aging despite preserved HNF4α expression. By integrating transcriptomic, epigenomic, chromatin occupancy, spatial, and functional approaches, we define an age-dependent HNF4α regulatory landscape and show that aging selectively disrupts HNF4α-dependent metabolic and identity programs while derepressing inflammatory and alternative cellular programs. Importantly, hepatocyte-specific loss of HNF4α recapitulated key features of liver aging, whereas restoration of HNF4α in aged liver reversed steatosis and reactivated critical metabolic programs. Together, these findings identify HNF4α as a transcriptional gatekeeper of hepatocyte state and supports HNF4α activity and its gene programs as a promising therapeutic target to preserve hepatocyte metabolic function and identity during aging.

Our findings reveal a disconnect between HNF4α abundance and transcriptional activity during aging. Despite preserved HNF4α expression, its chromatin occupancy and transcriptional output are reduced with age, indicating that transcription factor abundance alone may not accurately reflect functional activity in aged tissues. Consistent with this concept, Gunewardena et al. established that HNF4α activity, rather than expression, is a key determinant of liver disease severity and prognosis^19^. Our findings extend this observation to physiological aging and demonstrate that reduced HNF4α activity reflects remodeling of a broad regulatory network encompassing hepatocyte identity, lipid and intermediary metabolism, inflammatory signaling, and alternative cellular states.

The mechanisms underlying this uncoupling remain to be determined, but our data implicate remodeling of the regulatory chromatin environment. HNF4α-associated regulatory regions exhibited reduced H3K27ac, diminished promoter-enhancer interactions, reduced chromatin accessibility, and enrichment of HNF4α motifs at age-hypermethylated promoters. Most importantly, HNF4α CUT&RUN demonstrated reduced chromatin occupancy across the aged liver. These findings link age-associated chromatin remodeling with impaired engagement of a lineage-defining transcription factor. However, the relationship is unlikely to be strictly unidirectional. Reduced accessibility at HNF4α-associated regions may constrain HNF4α binding, while diminished HNF4α activity could further impair maintenance of active regulatory elements^29, 30^. Thus, additional studies are needed to delineate the regulatory link between age-dependent epigenetic remodeling and loss of HNF4α activity and to what degree these two phenomena are a linear cause-and-effect, act independently or form a mutually reinforcing cycle. Other mechanisms may also contribute to this uncoupling, including age-associated changes in HNF4α post-translational modification, interacting transcriptional partners or chromatin regulators, and the hepatic lipid and metabolite environment. In particular, changes in FA composition could influence HNF4α through its ligand-binding domain. Defining how these mechanisms interact will be important for determining whether restoring chromatin accessibility, HNF4α activity, or both is required to fully restore youthful hepatocyte function.

A major advance of this study is the experimental definition of HNF4α target genes in vivo. Integration of HNF4α occupancy with hepatocyte-specific genetic loss-of-function distinguished genes that are both directly associated with HNF4α chromatin occupancy and functionally dependent on HNF4α. This revealed that HNF4α regulates a broader network beyond its canonical role as an activator of hepatocyte metabolic genes. In young liver, HNF4α-dependent transcription comprised complementary programs that promote hepatocyte identity, lipid metabolism, and metabolic homeostasis while restraining inflammatory, stress, proliferative, developmental, and alternative cellular programs. HNF4α therefore appears to function as a transcriptional gatekeeper that simultaneously reinforces the differentiated hepatocyte state and constrains transcriptional programs that are incompatible with that state. The spatial organization of these targets across periportal, midlobular, and pericentral hepatocytes further indicates that HNF4α regulation is integrated with the functional specialization of the liver lobule. Age-associated disruption of these spatially organized programs may therefore compromise not only the magnitude of hepatocyte transcription but also its regional organization.

Importantly, aging did not produce a uniform loss of HNF4α regulation. HNF4α-activated targets involved in hepatocyte identity and core metabolic functions were preferentially lost with age, whereas HNF4α-inhibited targets involved in inflammatory signaling, adhesion, migration, and alternative cellular programs were preferentially induced. At the same time, a substantial fraction of HNF4α targets remained relatively stable, and a smaller subset changed in the opposite direction. Thus, aging appears to selectively rewire the HNF4α regulatory network, with some regulatory programs being more vulnerable than others. This distinction is important because it suggests that age-associated HNF4α dysfunction reflects selective remodeling of regulatory capacity rather than complete loss of HNF4α function.

The functional consequences of this remodeling were supported by characterization of hepatocyte-specific *Hnf4a* deletion in the liver. HNF4α deletion in young liver reproduced approximately 63% of age-associated transcriptional changes, including reduced hepatocyte metabolic programs and increased inflammatory and IFN-associated transcription. Notably, the high proportion of HNF4α-activated targets among concordantly age-downregulated genes (751/1,194; 62.9%) supports a prominent role for HNF4α in directly maintaining the core hepatocyte metabolic transcriptional programs that decline with age. Conversely, the relatively low proportion of HNF4α-inhibited targets among the age-upregulated genes (209/1,467; 14.2%) suggests a greater contribution of secondary indirect signaling cascades downstream of HNF4α for the broader induction of inflammatory and immune-associated programs. Furthermore, HNF4α deletion was sufficient to induce steatosis and immune infiltration in young mice and exacerbated the pre-existing steatohepatitis phenotype of aged liver, demonstrating that aged hepatocytes remain dependent on residual HNF4α activity despite its age-associated suppression. Thus, reduced HNF4α activity is unlikely to represent merely a consequence of hepatic aging but can reproduce key features of age-associated hepatic dysfunction and MASLD.

The selective disruption of HNF4α-dependent metabolic programs provides a potential mechanism linking reduced HNF4α activity to hepatic steatosis. Age-downregulated HNF4α-activated targets were enriched for FA oxidation, mitochondrial and peroxisomal metabolism, lipid handling, and turnover, suggesting diminished capacity to process and dispose of excess lipid. Conversely, selective induction of lipid-storage-associated genes, including Cidec and Soat2, may favor lipid sequestration and altered intracellular partitioning. Although both HNF4α regulatory classes encompass metabolic genes, their distinct functional composition suggests that aging disrupts the coordinated balance of lipid acquisition, utilization, and storage rather than uniformly suppressing lipid metabolism. These changes suggest liver aging may shift hepatic lipid homeostasis toward net accumulation.

Our findings directly implicate HNF4α dysfunction to hepatic inflammation and immune remodeling. Loss of HNF4α induced inflammatory and interferon-associated transcriptional programs, while HNF4α occupancy identified direct regulation of selected inflammatory pathway components, including *Ifngr1*. The broader ISG response is likely amplified through secondary signaling rather than being directly controlled by HNF4α. Moreover, age-associated derepression of HNF4α-inhibited genes involved in adhesion, vascular signaling, and extracellular matrix remodeling may alter the hepatic microenvironment and facilitate immune-cell recruitment. Consistent with this model, HNF4α deletion altered hepatic immune composition, reproducing selected age-associated changes while also inducing distinct responses. These observations suggest that HNF4α dysfunction may contribute to a reinforcing relationship between hepatocyte metabolic stress, inflammatory signaling, and immune remodeling, although it does not fully account for the complex immune changes accompanying physiological aging.

Perhaps most importantly, our restoration experiments demonstrate that aged hepatocytes retain substantial HNF4α-dependent regulatory capacity. Ectopic HNF4α expression reduced hepatic lipid accumulation, remodeled the lipidome, and reactivated age-suppressed genes involved in FA metabolism, mitochondrial function, and hepatocyte identity. These improvements occurred without complete restoration of the youthful transcriptome, indicating that recovery of critical metabolic functions does not require wholesale reversal of the aged transcriptional state. This residual plasticity is consistent with our observation that HNF4α-associated chromatin accessibility is reduced, but not eliminated, during aging. Interestingly, metabolic programs were preferentially restored, whereas inflammatory and interferon-associated programs remained incompletely resolved. This differential response may reflect the extensive direct regulation of hepatocyte metabolic genes by HNF4α, compared with inflammatory networks that can be sustained through secondary signaling and intercellular communication. Consequently, HNF4α restoration may be particularly effective in reversing hepatocyte-intrinsic metabolic dysfunction, while established inflammatory remodeling may require complementary interventions targeting the broader tissue environment.

Several limitations should be considered. First, although combined chromatin occupancy and genetic perturbation provide strong evidence for direct HNF4α regulation, they do not establish whether individual age-associated regulatory changes are caused by altered HNF4α activity, chromatin remodeling, changes in other transcriptional regulators, or altered hepatocyte state. Context-dependent regulation and compensatory activity of other transcription factors may also contribute to the selective remodeling of HNF4α targets. Second, although hepatocyte-targeted manipulation and spatial/single cell analyses strengthen the interpretation of hepatocyte-intrinsic regulation, aging-associated changes in cellular composition, intercellular signaling, and cell identity complicate distinctions between cell-autonomous from non-cell-autonomous mechanisms. Finally, HNF4α restoration was evaluated over a relatively short period and therefore establishes reversibility of selected age-associated phenotypes rather than complete reversal of liver aging.

In summary, our findings establish HNF4α as a critical regulator of hepatocyte identity and metabolic homeostasis whose transcriptional activity becomes impaired during aging despite preserved expression. Age-associated remodeling of the chromatin landscape is associated with reduced HNF4α occupancy and selective disruption of HNF4α-dependent programs, shifting the balance from differentiated metabolic function toward lipid accumulation and inflammatory and alternative cellular states. Genetic loss of HNF4α recapitulates key features of hepatic aging, while restoration of HNF4α reverses steatosis and restores critical metabolic programs. These findings support a model in which HNF4α is impaired but not absent during aging and identify residual HNF4α-dependent regulatory capacity as a potentially targetable source of hepatic plasticity. More broadly, they suggest that targeting lineage-defining transcription factors may provide a strategy for reversing selected aspects of tissue aging even when the abundance of the transcription factor itself is preserved.

## Materials & Methods

### Animals and experimental design

Young adult (4 month) and aged (21 month) C57BL/6J male mice were obtained from the National Institute on Aging (NIA) Aged Rodent Colony through Charles River Laboratories. Mice were maintained at the Sanford Burnham Prebys Medical Discovery Institute animal facility under controlled temperature (22.5°C) and a 12-h light/dark cycle, with ad libitum access to water and standard chow (Teklad Global 18% Protein Rodent Diet, Envigo, #2018). Animals were housed at up to five mice per cage. All animal procedures were performed in accordance with protocols approved by the Sanford Burnham Prebys Institutional Animal Care and Use Committee (IACUC) and institutional animal welfare guidelines. Mice were euthanized by CO₂ inhalation immediately before tissue collection in accordance with institutional guidelines.

For hepatocyte-specific *Hnf4a* deletion, young and aged mice received adeno-associated virus serotype 8 (AAV8) encoding Staphylococcus aureus Cas9 (SaCas9) and a single-guide RNA (sgRNA) targeting Hnf4a, with an sgRNA targeting the Rosa26 locus serving as the control. For HNF4α overexpression, young and aged mice received AAV8 encoding mouse HNF4α or a fluorescent reporter as control. AAV8 vectors were administered by retroorbital injection suspended in 100 μL sterile PBS per mouse at a dose of 3.33 x 10^11^ viral genomes for KO and 1 x 10^11^ viral genomes for overexpression studies. Livers were collected three weeks after administration for molecular and histological analyses.

### AAV vector construction and production

AAV expression vectors were generated using plasmids obtained from VectorBuilder, including pAAV[Exp]-TBG>mApple:WPRE and pAAV[Exp]-TBG-mHnf4a[NM_008261.3]/3xFLAG:P2A:mApple:WPRE. The CRISPR-Cas9 vector was derived from pX602-AAV-TBG::NLS-SaCas9-NLS-HA-OLLAS-bGHpA;U6::BsaI-sgRNA (Addgene #61593). Guide RNAs targeting mouse *Hnf4a* and *Rosa26* were designed using CRISPick with the GRCm38 mouse reference genome (Ensembl v102). sgRNA sequences were cloned into the pX602 AAV vector and transformed into NEB Stable competent Escherichia coli. Successful guide insertion was verified by Sanger sequencing using the universal U6 primer (5′-GACTATCATATGCTTACCGT-3′). Recombinant AAV8 particles were produced by the Sanford Burnham Prebys Functional Genomics Core Facility as previously described^26^. Briefly, HEK293T cells were cotransfected with the AAV expression vector, RepCap8, and helper plasmids (Cell Biolabs) using polyethylenimine (Polysciences). Viral particles were harvested 96 h after transfection from culture supernatants by polyethylene glycol precipitation and from cell pellets by sonication. Following benzonase treatment, viral particles were purified by iodixanol gradient ultracentrifugation, buffer-exchanged into phosphate-buffered saline (PBS), aliquoted, and stored at −80°C. Viral titers were determined by SYBR Green-based quantitative PCR.

### Validation of CRISPR-mediated *Hnf4a* deletion

Genomic DNA was isolated from liver tissue using the Monarch Genomic DNA Purification Kit (New England Biolabs, #T3010S). Regions surrounding the sgRNA target sites were amplified by PCR using 2× Taq PCR Master Mix (APExBIO, #K1034) and gene-specific primers. PCR products were purified using the QIAquick PCR Purification Kit (QIAGEN, #28104) and subjected to Sanger sequencing (Genewiz). Editing efficiency was evaluated using Inference of CRISPR Edits (ICE) analysis, version 3.0 (Synthego), as previously described^26^. *Hnf4a* deletion was further validated by Western blot analysis of HNF4α protein abundance.

### Hepatocyte isolation and whole liver tissue collection

Primary hepatocytes were isolated from young and aged mouse livers using a previously published two-step perfusion protocol^26^. Briefly, a 24-gauge catheter was inserted into the inferior vena cava, and livers were perfused with prewarmed Hank’s balanced salt solution (HBSS; Gibco, #14175-095) containing EDTA, followed by digestion with Dulbecco’s modified Eagle medium (DMEM; Gibco, #10313-021) supplemented with 5 mM CaCl₂ and 1.2 mg/mL type IV collagenase (Gibco, #17104-019). Following perfusion, livers were excised and dissociated, and hepatocytes were enriched by five sequential low-speed centrifugations (50 x g, 2 min) in ice-cold DMEM containing 2% bovine serum albumin. Hepatocytes were further purified using a 90% Percoll density gradient. Isolated hepatocytes were processed for downstream molecular analyses as previously described^26^. For whole-liver analyses, tissues were collected immediately following euthanasia and processed according to the requirements of each assay. Samples for molecular profiling were snap-frozen in liquid nitrogen and stored at −80°C. Tissues for histological analyses were either formalin-fixed and stored at 4°C (Immunohistochemistry) or frozen embedded in OCT and stored at −80°C (Oil Red O Staining and MERFISH).

### RNA isolation and quantitative PCR

Total RNA was extracted from mouse liver using TRIzol reagent (Invitrogen, #15596018) according to the manufacturer’s instructions, with modifications to the phenol-chloroform extraction procedure. RNA concentration and purity were assessed using a NanoDrop One spectrophotometer (Thermo Fisher Scientific). For reverse transcription quantitative PCR (RT-qPCR), RNA was converted to complementary DNA using RevertAid Reverse Transcriptase (Thermo Fisher Scientific, #EP0441), RiboLock RNase Inhibitor (#EO0381), and the supplied reaction buffer. Quantitative PCR was performed using PowerUp SYBR Green Master Mix (Applied Biosystems, #A25741) on a QuantStudio 6 Flex Real-Time PCR System (Applied Biosystems, #4485691). Relative transcript abundance was calculated using ΔΔCt method and normalized to *Gapdh* and *Hprt* as housekeeper reference genes.

### Bulk RNA sequencing and differential gene expression analysis

Bulk RNA sequencing was performed on isolated hepatocytes from young and aged wild-type (WT) mice and on whole-liver tissues collected following hepatocyte-specific *Hnf4a* deletion or overexpression. Previously published hepatocyte RNA-seq data from isolated young and aged mice were generated as previously described^26^ and reanalyzed for the present study to investigate age-associated HNF4α-dependent transcriptional programs. For newly generated whole-liver RNA-seq datasets, RNA quality was assessed using an Agilent TapeStation before library preparation and sequencing by the Sanford Burnham Prebys Genomics Core Facility. Poly(A) RNA-seq libraries were prepared using an epMotion 5075 automated liquid handling system (Eppendorf), according to the library preparation manufacturer’s protocol and sequenced using paired-end reads on an Element Biosciences AVITI system. Raw sequencing reads were processed by the Sanford Burnham Prebys Bioinformatics Core Facility using the nf-core/rnaseq pipeline (v3.14.0), implemented in Nextflow, with the parameters --aligner star_rsem --igenomes_ignore --genome null --gencode. Reads were aligned to the mouse GRCm38 primary genome assembly using STAR, and transcript abundance was quantified using RSEM with GENCODE release M25 annotations. Differential expression analysis was performed using DESeq2. Genes with fewer than 10 total raw counts across all samples were excluded before differential expression analysis. Statistical significance was assessed using Wald tests with Benjamini-Hochberg false discovery rate (FDR) correction. Differentially expressed (DE) genes were identified using an adjusted p-value threshold of ≤ 0.05, with additional fold change (FC) criteria applied as indicated for individual analyses.

### Lipidomic profiling and analysis

Targeted lipidomic profiling was performed on lipid extracts of whole-liver samples from PBS-perfused WT young and aged mice or from unperfused mice receiving AAV8-mediated HNF4α overexpression or control vectors. Liver tissue was collected and submitted to the Functional Lipidomics Core at the Barshop Institute, UT Health San Antonio, for lipid extraction, mass spectrometry, and lipid identification. Lipid species were analyzed using multidimensional mass spectrometry-based shotgun lipidomic analysis^31^. In brief, liver tissue homogenate containing 1.0 mg of protein which was determined with a Pierce BCA assay was accurately transferred to a disposable glass culture test tube. A premixture of lipid internal standards (IS) species, including D14:1 choline glycerophospholipid (PC), D16:1 ethanolamine glycerophospholipid (PE), D15:0 phosphatidylglycerol (Na) (PG), D14:0 phosphatidylserine (Na) (PS), D14:0 phosphatidic acid (Na) (PA), D4-16:0 free fatty acid (FFA), N12:0 sphingomyelin (SM), 14:0 Lyso ethanolamine glycerophospholipid (LPE), 17:0 lyso choline glycerophospholipid (LPC), T14:0 cardiolipin (CL), N17:0 ceramide (Cer), 13C4 16:0 acyl carnitine (AC) and T17:1 triglyceride (TAG) (14.77,14.83, 2.06, 0.64, 0.39, 2.91, 0.82, 0.42, 1.03, 3.65, 0.72, 0.006 and 15.01 nmol/mg protein, respectively), was added prior to conducting lipid extraction for quantification of the targeted lipid species. Lipid extraction was performed using a modified Bligh and Dyer procedure^31^, and each lipid extract was reconstituted in chloroform:methanol (1:1, v:v) at a volume of 400 µL/mg protein. Phosphoethanolamine (PE) and free fatty acid (FFA) were derivatized as described previously^32, 33^ before lipidomic analysis. All internal standards were purchased from Avanti Polar Lipids (USA). For shotgun lipidomics, lipid extract was further diluted to a final concentration of ∼500 fmol total lipids per µL. Mass spectrometric analysis was performed on a triple quadrupole mass spectrometer (TSQ Altis, Thermo Fisher Scientific, San Jose, CA) and a Q Exactive mass spectrometer (Thermo Scientific, San Jose, CA), both of which were equipped with an automated nanospray device (TriVersa NanoMate, Advion Bioscience Ltd., Ithaca, NY) as described^34^. Identification and quantification of lipid species were performed using an automated software program^35, 36^. Data processing (e.g., ion peak selection, baseline correction, data transfer, peak intensity comparison and quantitation) was performed as described^36^. The result was normalized to the protein content (nmol lipid/mg protein). Differential lipid abundance was assessed between young and aged livers and between HNF4α-overexpressing and age-matched control livers. Lipid species with p < 0.05 were considered significantly different, as indicated in the figure legends. Lipidomic pathway enrichment was performed using MetaboAnalyst. The heatmap for differential lipid species was generated using Broad Institute Morpheus software. All other graphs and heatmaps were generated using GraphPad Prism software.

### Metabolomic profiling and analysis

Previously published untargeted metabolomic data from young and aged mouse livers were reanalyzed for the present study and processed as previously described^26^. Briefly, frozen liver tissues were extracted in ice-cold methanol:acetonitrile (5:3:2, v/v/v) at a concentration of 15 mg tissue/mL. Extracts were vortexed for 30 min at 4°C and centrifuged at 18,000 × g for 10 min at 4°C. Metabolites were separated using a Thermo Vanquish ultra-high-performance liquid chromatography system equipped with a Kinetex C18 column (150 x 2.1 mm, 1.7 μm; Phenomenex) and analyzed using a Thermo Orbitrap Exploris 120 mass spectrometer in positive and negative electrospray ionization modes. Mass spectra were acquired over an m/z range of 65-900. Raw files were converted to mzXML format using RawConverter, and metabolites were annotated and quantified using El-Maven (Elucidata), the KEGG database, and an in-house reference library of more than 600 compounds, as previously described^26^. Metabolomic data were log10-transformed and autoscaled by mean-centering and standard deviation. Metabolites with p < 0.05 were considered differentially abundant. Integrated pathway analyses of age-associated metabolomic and transcriptomic changes were performed using MetaboAnalyst v5.0 and KEGG pathway annotations, with FDR < 0.05 used to identify significantly enriched pathways.

### H3K27ac ChIP-seq analysis

Previously generated H3K27ac chromatin immunoprecipitation sequencing (ChIP-seq) data from isolated young and aged mouse hepatocytes were reanalyzed to examine age-associated changes in active chromatin associated with HNF4α-regulated genes. ChIP-seq was performed as previously described^26^. Briefly, hepatocyte chromatin was fragmented by sonication to approximately 250-500 bp using a Bioruptor Pico. Chromatin IP was performed using Protein G Dynabeads (Invitrogen, #1000D) with antibodies against histone H3 (Abcam, #ab1791) or H3K27ac (Abcam, #ab4729). Following overnight immunoprecipitation at 4°C, DNA was purified by phenol-chloroform extraction and ethanol precipitation and submitted for library preparation and sequencing. Sequencing reads were quality-checked using FastQC v0.11.8 and trimmed using Trim Galore v0.4.4 when necessary. Reads were aligned to a combined mm10/hg38 reference genome using STAR v2.5.3a to account for human spike-in DNA. Reads aligning to hg38 were excluded from downstream mouse genomic analyses. Duplicate reads were removed using Picard v2.18.4, and peaks were identified using HOMER findPeaks relative to matched input controls. Normalized H3K27ac signal was evaluated at genomic regions associated with IPA-defined HNF4α-regulated genes. Metagene profiles were generated using metagene2 v1.12.0, and ChIP-seq coverage was normalized using a log_2_-ratio normalization method. Age-associated changes in H3K27ac signal intensity were correlated with corresponding changes in hepatocyte gene expression using Pearson’s correlation coefficient (r), with statistical significance assessed using a two-tailed test in GraphPad Prism v11.0.2.

### H3K27ac HiChIP analysis

Previously generated H3K27ac HiChIP data from young and aged mouse hepatocytes^37^ were analyzed to evaluate age-associated changes in chromatin interactions associated with HNF4α-regulated genes. HiChIP data were generated and initially processed as described previously^37^. To facilitate comparisons between young and aged samples, chromatin contact reads were downsampled to an equal number per sample to control for differences in sequencing depth. Promoter-enhancer interactions associated with IPA-defined HNF4α target genes were identified from the processed HiChIP contact data. The number and strength of promoter-enhancer contacts associated with IPA-defined HNF4α target genes was compared between young and aged hepatocytes to assess age-associated changes in regulatory chromatin interactions.

### Bulk ATAC-seq and chromatin accessibility analysis

Previously generated bulk assay for transposase-accessible chromatin sequencing (ATAC-seq) data from young and aged male mouse livers (GEO: GSE243630) were reanalyzed to investigate age-associated changes in chromatin accessibility and predicted HNF4α occupancy at regulatory regions. Original sequencing data were processed as described in the GEO record. Accessible chromatin peaks were identified using MACS2 and classified as young-specific, aged-specific, or shared based on overlap between age-specific peak sets. Transcription factor occupancy prediction and motif-associated accessibility analyses were performed using TOBIAS, including BINDetect to identify HNF4α motif-containing regions and evaluate age-associated differences in predicted transcription factor occupancy. HNF4α motif enrichment in young-specific ATAC-seq peaks was evaluated relative to random genomic sequences and aged-specific peak sequences. Motif-containing accessible regions were annotated according to genomic location and associated genes using the mm10 mouse reference genome.

### Single-nucleus chromatin accessibility analysis

Single-nucleus chromatin accessibility data (snATAC-seq) from young (4 months), aged (23 months), and geriatric (30 months) male and female mouse livers were analyzed to identify transcription factor-associated regulatory regions exhibiting progressive age-associated loss of accessibility. snATAC-seq data were generated from 10x Genomics sn multiome and processed through the NIH Cellular Senescence Network (SenNet). Data are available through the SenNet Data Portal (https://doi.org/10.60586/SNT987.ZGSS.452). Age-associated chromatin accessibility changes were analyzed at the pseudobulk level using edgeR. Fragment counts within hepatocyte-accessible peaks were aggregated by biological replicate, and low-abundance peaks were filtered using filterByExpr (min.count = 10). Libraries were normalized using the trimmed mean of M-values (TMM) method. To identify progressive accessibility changes across the aging trajectory, age was modeled as a continuous covariate in an edgeR generalized linear model, and the age-associated coefficient was evaluated using a quasi-likelihood F-test (glmQLFit and glmQLFTest). Peaks exhibiting a significant negative age coefficient (Benjamini-Hochberg false discovery rate [FDR] < 0.05) were classified as regions of progressive age-associated accessibility loss. Transcription factor motif enrichment analysis was performed on these regions using HOMER (v5.1; findMotifsGenome.pl) with the mm10 mouse reference genome. Known-motif enrichment was assessed against the background of all peaks included in the differential accessibility analysis, retaining the original peak widths (-size given). Motifs with an adjusted q-value < 0.05 were considered significantly enriched.

### Whole-genome bisulfite sequencing and DNA methylation analysis

Whole-genome bisulfite sequencing (WGBS) was performed on primary hepatocytes isolated from young (5 months) and aged (22 months) C57BL/6 mice. Genomic DNA was extracted using the Quick-DNA Miniprep Kit (Zymo Research, #D3024), and DNA integrity was assessed using an Agilent TapeStation. Library preparation and bisulfite conversion were performed by BGI according to SOP-SS-023, version A0, followed by sequencing on an Illumina NovaSeq system. Raw sequencing reads were aligned using Bismark v0.15.0, and complementary CpG dyads were collapsed as previously described^13^. Differentially methylated regions (DMRs) were identified using methylKit pipeline, wherein significantly differentially methylated CpGs between conditions were called with a mean methylation difference above 25% and an FDR-corrected p-value below 0.001. Age-associated hypermethylated promoter regions were subjected to transcription factor motif enrichment analysis using HOMER with the mm10 mouse reference genome to identify candidate transcription factors associated with increased promoter methylation during aging.

### HNF4α CUT&RUN and chromatin occupancy analysis

HNF4α chromatin occupancy was assessed in young and aged mouse livers (n = 4 per age group) using the ChIC/CUT&RUN Assay Kit (Active Motif, #53180). Briefly, nuclei were isolated from frozen liver tissue using the gentleMACS Dissociator system (Miltenyi Biotec) according to the manufacturer’s instructions. CUT&RUN was performed using antibodies against HNF4α (Invitrogen, #MA1-199), H3K4me3 (Active Motif, #39916) as a positive control, and IgG (Active Motif, #106138) as a negative control, following the manufacturer’s protocol. CUT&RUN libraries were prepared by the Sanford Burnham Prebys Genomics Core using an epMotion 5075 automated liquid handling system (Eppendorf) and sequenced on an Element Biosciences AVITI platform. Raw sequencing reads were aligned to the mm10/GRCm38 mouse reference genome using Bowtie2. Aligned reads were sorted and indexed using SAMtools, and duplicate reads were identified and removed using Picard MarkDuplicates. For each sample, IgG controls were used for background normalization. HNF4α binding peaks were called using MACS2. Consensus HNF4α binding peaks were independently defined for young and aged livers from narrowPeak files, retaining peaks detected in at least three of four biological replicates within each age group. Consensus peaks were annotated according to genomic location, including promoter/transcription start site regions, gene bodies, and intergenic regions, using HOMER with the mm10 mouse reference genome. Age-associated differences in HNF4α occupancy were evaluated by comparing peak numbers and genomic distributions between young and aged livers. For visualization, CUT&RUN coverage tracks were normalized using bins-per-million (BPM) normalization, and average normalized coverage was calculated across biological samples within each condition. Age-specific consensus peak sets were subsequently integrated with corresponding *Hnf4a* knockout RNA-seq data to identify HNF4α-dependent target genes, as described below.

### MERFISH spatial transcriptomic analysis

Multiplexed error-robust fluorescence in situ hybridization (MERFISH) was performed using the MERSCOPE platform (Vizgen) to characterize spatially resolved transcriptional programs in young and aged mouse livers. Spatial transcriptomic profiling was performed using a custom 500-gene panel containing markers of hepatic cell identity, zonation, and age-associated transcriptional responses, including 126 experimentally defined HNF4α target genes. MERFISH data were generated from two young and two aged mouse livers by Sanford Burnham Histology and Genomic Cores per manufacturer’s instructions. MERSCOPE images were segmented using the Vizgen Post-processing Tool (VPT, v1.3.0) with Cellpose (2D cyto2 and nuclei models). DAPI and PolyT image channels were normalized using contrast-limited adaptive histogram equalization (CLAHE) before segmentation. Cell and nuclear masks were generated from the central z-plane (z = 3) and combined into final cell boundaries using default VPT fusion parameters. Cells containing more than 100 transcripts and more than 10 unique genes were retained for downstream analyses. MERFISH expression data were SCTransformed, integrated using Harmony, and clustered by uniform manifold approximation and projection (UMAP) in Seurat v5.0.1.9001. Individual clusters were annotated using a panel of liver cell-type marker genes and the top 20 cluster-specific genes evaluated using CellKB. Hepatocytes were classified into periportal, midlobular, and pericentral populations based on their transcriptional profiles and established zonation markers included in the cell-type annotation panel. For spatial differential expression analyses, pseudobulk gene counts were generated by summing transcript counts across cells belonging to each hepatocyte zonal population within each biological replicate. Pseudobulk counts were normalized and analyzed for differential expression using DESeq2 v1.30.0. Age-associated differential expression was evaluated independently within periportal, midlobular, and pericentral hepatocyte populations. Genes with a Benjamini-Hochberg false discovery rate (FDR)-adjusted p-value ≤ 0.05 were considered significantly DE. Zonal enrichment of HNF4α target genes in young liver was assessed by comparing expression across periportal, midlobular, and pericentral hepatocyte populations. Genes exhibiting significant age-associated changes in one or more hepatocyte zones were classified according to the direction and spatial distribution of their expression changes. Overlap among zonal differential expression sets was visualized using UpSet plots. Spatial expression maps were generated for selected periportal and pericentral HNF4α target genes, including Cyp2f2, Gls2, Glul, and Nr1i3.

### Identification and classification of HNF4α-dependent target genes

High-confidence HNF4α-dependent target genes were identified by integrating HNF4α CUT&RUN chromatin occupancy with differential gene expression following hepatocyte-specific Hnf4a deletion. Young and aged HNF4α target sets were independently defined using age-matched CUT&RUN consensus peaks and whole-liver RNA-seq comparisons between Hnf4a knockout and Rosa26 control mice. Genes were retained as HNF4α targets if they exhibited significant differential expression following Hnf4a deletion (FDR < 0.05) and were associated with at least one HNF4α CUT&RUN peak within the gene body, promoter, or an intergenic region within 5 kb of the transcription start site. HNF4α target genes were further classified according to their transcriptional response to Hnf4a deletion. Genes downregulated following deletion were designated HNF4α-activated targets, whereas genes upregulated following deletion were designated HNF4α-inhibited targets. Target genes were independently classified in young and aged liver, and the resulting gene sets were compared to identify shared and age-specific HNF4α-dependent regulatory networks.

### Functional enrichment and pathway analysis

Functional enrichment analyses were performed using STRING (Search Tool for the Retrieval of Interacting Genes/Proteins; https://string-db.org/), Ingenuity Pathway Analysis (IPA; QIAGEN), WebGestaltR v0.4.4, MetaboAnalyst v5.0, including gene sets from Gene Ontology Biological Process (GO:BP), Reactome, Hallmark, and KEGG, as appropriate. IPA upstream regulator analysis was performed on age-associated DE hepatocyte genes to identify candidate transcriptional regulators and predict their activation or inhibition based on the direction of target gene expression changes. IPA-defined HNF4α-regulated genes were extracted for downstream transcriptional and epigenomic integration. Gene set over-representation analysis and gene set enrichment analysis were performed using STRING v12.5 and WebGestaltR v0.4.4. Functional enrichment results were ranked and visualized using adjusted p-values, enrichment ratios, gene counts, or enrichment scores, as indicated in the figures.

### Age-associated remodeling of HNF4α target gene networks

To determine how HNF4α-dependent transcriptional programs change during physiological aging, experimentally defined HNF4α target gene sets were integrated with previously published differential gene expression data from isolated young and aged hepatocytes. Young- and aged-defined HNF4α-activated and HNF4α-inhibited target genes were evaluated for enrichment among genes upregulated or downregulated with age. Young-defined HNF4α target genes were further stratified according to their age-associated expression patterns into age-upregulated, age-downregulated, or not significantly changed categories. Functional enrichment was performed independently for each category to identify biological processes associated with age-sensitive and age-intact HNF4α-dependent transcriptional programs. Age-associated changes in HNF4α target gene expression were also examined in spatial MERFISH data to evaluate the extent to which transcriptional remodeling was shared across hepatocyte zones or restricted to specific zonal populations.

### Transcriptomic comparisons of physiological aging and HNF4α perturbation

To assess the extent to which hepatocyte-specific HNF4α loss recapitulates age-associated transcriptional changes, differential expression results from young Hnf4a knockout versus young Rosa26 control livers were compared with those from aged versus young WT hepatocytes. Genes were matched across datasets, and log2FC was compared using Pearson correlation and simple linear regression, with statistical significance assessed using an F-test. Genes significantly changed in both comparisons were classified as concordantly upregulated, concordantly downregulated, or discordantly regulated based on the direction of their expression changes. To evaluate transcriptional responses to HNF4α restoration, differential expression following AAV8-mediated HNF4α overexpression in aged livers was compared with age-associated transcriptional changes in WT hepatocytes. Genes displaying opposing expression changes between aging and HNF4α restoration were identified and subjected to functional enrichment analysis.

### Western blot analysis

Frozen mouse liver tissues were homogenized using a Precellys tissue homogenizer in RIPA lysis buffer with 1x protease and phosphatase inhibitors. Protein concentrations were determined using Bradford reagent (Pierce, #1863028) and a SpectraMax 190 plate reader. Protein lysates were separated by SDS-PAGE using Bio-Rad Mini-PROTEAN precast gels and transferred to nitrocellulose membranes using a Bio-Rad Trans-Blot Turbo transfer system. Membranes were stained with Ponceau S (Sigma-Aldrich, #P7170) to assess protein loading and transfer efficiency, followed by blocking in 5% nonfat milk in Tris-buffered saline containing Tween-20 (TBST) for 1 h at room temperature. Membranes were incubated overnight at 4°C with rabbit anti-HNF4α antibody (Abcam, #ab181604), followed by goat anti-rabbit IgG-HRP secondary antibody (Millipore, #AP307P; RRID) for 2 h at room temperature. Immunoreactive proteins were detected using SuperSignal West Pico PLUS chemiluminescent substrate and imaged using a Bio-Rad ChemiDoc Touch imaging system. Band intensities were quantified using ImageJ software and normalized to Ponceau S total protein staining.

### Histological assessments and quantifications

Hepatic lipid accumulation was assessed by Oil Red O staining (ORO) from OCT-embedded frozen liver sections. Frozen tissues were cryosectioned at 10 μm thickness, fixed in 10% neutral-buffered formalin for 10 min, washed, and stained according to the manufacturer’s protocol (Poly Scientific, Cat. #K043). Immunohistochemical (IHC) staining was performed on formalin-fixed, paraffin-embedded (FFPE) liver sections to assess hepatic immune-cell populations, including CD45⁺ leukocytes, F4/80⁺ macrophages/Kupffer cells, B220⁺ B cells, and CD3⁺ T cells. Liver tissues were fixed in 10% neutral-buffered formalin (Epredia, Cat. #9400-1) for 24 h at room temperature, paraffin-embedded, and sectioned at 5 μm thickness. Sections were deparaffinized and rehydrated through xylene and graded ethanol solutions, followed by heat-induced antigen retrieval for 40 min using a heated steamer. Primary antibodies included anti-CD45 (1:500; Cell Signaling Technology, Cat. #70257S; pH 6 retrieval), anti-F4/80 (1:300; Cell Signaling Technology, Cat. #70076S; pH 6 retrieval), anti-CD3 (1:200; Abcam, Cat. #ab16669; pH 9 retrieval) and anti0B220 (1:100; Invitrogen, Cat. #14-0452; pH 6 retrieval). Sections were incubated with primary antibodies overnight, followed by chromogenic detection using 3,3′-diaminobenzidine (DAB) and hematoxylin counterstaining. All stained slides were scanned at 20x magnification using a Leica Aperio AT2 digital slide scanner, and whole-slide images were visualized using Aperio ImageScope software (Leica Biosystems). Lipid droplet accumulation was quantified using the OpenCV library for red droplet isolation via HSV color segmentation, followed by a 2 x 2 pixel morphological opening to refine droplet boundaries. Droplet count, size, and total area were quantified using skimage.measure. Digital images of IHC whole tissue scans were subjected to ChatGPT-assisted computational image analysis using a color-based approach to distinguish brown 3,3′-diaminobenzidine (DAB)-positive staining from hematoxylin counterstaining and background. Tissue regions were segmented to exclude non-tissue background, and DAB-positive pixels were identified using stain-specific intensity thresholds. Positive staining was quantified as the percentage of analyzed tissue area occupied by DAB-positive pixels and expressed as the percentage of tissue area occupied by positive staining.

### Flow cytometry

Liver immune-cell populations were characterized by flow cytometry following hepatocyte-specific *Hnf4a* deletion in young and aged mice. Fresh liver L lobe tissues were collected in ice-cold RPMI 1640 supplemented with 10% FBS and mechanically dissociated through a 70-μm cell strainer and rinsed with 20 mL complete RPMI. Hepatocytes were removed by low-speed centrifugation (60 x g, 2 min, 4°C, no brake). The supernatant was collected and centrifuged at 420 x g for 4 min at 4°C with brake on. Cell pellets were resuspended in 10mL 40% Percoll (Cytiva) in HBSS and centrifuged at 850 x g for 25 min at 20°C without brake to enrich nonparenchymal immune cells. After the supernatant was discarded, residual erythrocytes were lysed in 1 mL ACK lysis buffer (KD Medical) for 3 min at room temperature. Lysis was stopped with 2 mL complete RPMI, and cells were centrifuged at 420 x g for 4 min at 4°C and resuspended in 5 mL complete RPMI. Viable cells were counted using a hemocytometer, transferred to 96-well U-bottom plates, and resuspended in FACS buffer (PBS containing 2% FBS). Cell suspensions were stained with LIVE/DEAD Fixable Red viability dye (Invitrogen, #L34972; 1:1,000) for 15 min at room temperature, followed by Fc receptor blockade with anti-CD16/32 antibody (BioLegend; 1:500). Cells were stained with fluorophore-conjugated surface antibodies for 30 min at 4°C. For intracellular FoxP3 detection, cells were fixed and permeabilized using the FoxP3 Transcription Factor Staining Buffer Set (eBioscience) according to the manufacturer’s instructions and then stained with intracellular antibodies for 45 min at 4°C. Samples were acquired on a BD FACSymphony A3 cell analyzer using FACSDiva v9.1 and analyzed using FlowJo v10.10.0 (BD Biosciences). Immune-cell populations were identified using separate T-cell and myeloid staining panels (Supp. Table 1 & 2). Single-color compensation controls and unstained controls were acquired for both panels; fluorescence-minus-one controls were included for selected T cell markers. T cell events were gated sequentially by lymphocyte light scatter, two singlet gates, viabil ity, and CD45 positivity, followed by CD4 or CD8 expression. Regulatory T cells (Tregs) were identified as CD4⁺FoxP3⁺ and conventional CD4 T cells as CD4⁺FoxP3⁻. In the myeloid panel, two singlet gates and viability preceded the CD45hi gate; CD3ε⁺ T cells and B220⁺ B cells were identified in the CD11b-lymphocyte branch, and F4/80⁺ macrophages within the CD11b⁺MHC-IIlow/-Ly6Clow/-branch. Population frequencies shown in Supp. Fig. 4I-N were reported relative to the CD45hi immune-cell compartment. Antibody clones, fluorophore conjugates, manufacturers, catalog numbers, and working dilutions are provided in Supplementary Table 1 & 2.

### Statistical analysis

Statistical analyses were performed using R v≥4.0.0, GraphPad Prism v11.0.2, and Microsoft Excel, as appropriate. Biological replicates represent individual mice. Statistical tests were selected according to the experimental design and data characteristics, with specific tests, sample sizes, and significance thresholds provided in the corresponding Methods sections and figure legends. For comparisons involving multiple experimental groups, appropriate post hoc tests and multiple-comparison corrections were applied as indicated in the figure legends. Gene set and pathway enrichment analyses based on overrepresentation were performed using Fisher’s exact test, with the background gene universe defined according to the corresponding analysis. Correlations between continuous variables were assessed using Pearson’s correlation coefficient (r). For simple linear regression analyses, statistical significance was evaluated using an F-test. Multiple-testing corrections were applied to transcriptomic, epigenomic, and other high-dimensional analyses as described in the corresponding Methods sections. Data are presented as mean ± SEM, mean, unless otherwise indicated. A nominal p-value < 0.05 was considered statistically significant for analyses without multiple-testing correction. Adjusted p-values or false discovery rate (FDR) thresholds were used for high-dimensional analyses as specified in the corresponding Methods sections. Sample sizes, statistical tests, and significance thresholds are reported in the figure legends.

## Data availability

Previously published and newly generated sequencing datasets analyzed in this study are available through the NCBI Gene Expression Omnibus (GEO) and the NIH SenNet Data Portal. Relevant datasets include Hepatocyte RNAseq: GSE313646, bulk ATAC-seq (GSE243630), H3K27ac ChIP-seq (GSE279751), H3K27ac HiChIP (GSE253617), whole-genome bisulfite sequencing (GSE313655), HNF4α CUT&RUN (GSE338169), Hnf4a knockout RNA-seq (GSE338093), Hnf4a overexpression RNA-seq (GSE338094), and MERFISH spatial transcriptomics (GSE313704). Additional single-nucleus data are available through the SenNet Data Portal (https://doi.org/10.60586/SNT987.ZGSS.452).

## Acknowledgements

We thank the SBP Animal Facilities Core, especially Buddy Charbono and Adriana Charbono for their excellent technical assistance in AAV administration and animal care, respectively. We also thank Chun-Teng Huang in the SBP Functional Genomics Core for exceptional AAV production. Finally, tremendous thanks to the Nathan Shock Center and Functional Lipidomics Core at the Barshop Institute, UT Health San Antonio for help with planning, execution, analysis and interpretation of lipidomics data.

## Author Contributions

Conceptualization: JP, PDA. Methodology: JP, AH, MGT, AR, HT, MAL, LW, AG, RA, AD, AL, GG, MS, RP, AD’A, XH. Data generation and analysis: JP, KL, XL, AH, AR, MAL, GE, SM, LW, RM, AD’A, XH. Writing: JP, KL, PDA. Funding: JP, PDA. Supervision: SLB, AW, KYY, SK, XH, AEW, PDA

## Supplementary Figure Legends

**Supplementary Figure 1. Age-associated liver lipidomic and metabolic dysfunction is associated with epigenetic remodeling and repression of HNF4α transcriptome. A**) Quantification of lipid class abundances in young (Y) and old (O) mouse livers, normalized to total protein content (nmol/mg protein). **B**) Heatmap of differentially abundant lipids between Y and O livers. **C**) Integrated pathway enrichment analysis of O vs Y liver lipidomic and hepatocyte transcriptomic datasets using MetaboAnalyst. **D**) Scatterplot of gene length (log10 scale) versus mean normalized expression (log10 BaseMean) for all expressed genes in hepatocyte RNA-seq, highlighting IPA-defined HNF4α target genes downregulated with age (blue). Dashed lines indicate the median gene length (orange) and median expression (green) across expressed genes. Percentages indicate the proportion of highlighted IPA-HNF4α target genes within respective quadrants. Of the highlighted HNF4α targets, 57.4% were above the median gene length and 36.5% were below, with 93.9% exhibiting expression above the median hepatocyte gene expression. **E**) Reactome pathway enrichment analysis of 328 IPA-defined downregulated HNF4α target genes, as highlighted in D. **F**) Violin plots of hepatocyte *Hnf4a* RNA expression in Y and O mice using the Tabula Muris Senis dataset. **G**) H3K27ac HiChIP analysis of previously generated Yand O isolated mouse hepatocyte datasets, showing the number of associated enhancers, total promoter-enhancer contacts, and cumulative contact strength for IPA-defined HNF4α target genes. **H**) Genomic distribution of consensus HNF4α CUT&RUN peaks in young and aged mouse livers, classified by promoter/transcription start site, enhancer, gene body, and intergenic regions.

**Supplementary Figure 2. Integrated HNF4α chromatin occupancy and hepatocyte-specific genetic perturbation define high-confidence HNF4α target gene networks in young and aged liver. A**) Representative Sanger sequencing chromatograms of the *Hnf4a* sgRNA-targeted genomic region in control and hepatocyte-specific *Hnf4a* knockout (KO) in Y and O livers, illustrating sequence alterations following CRISPR-mediated *Hnf4a* editing. **B-C**) Hallmark pathway enrichment analysis of **B**) Y-HNF4α-activated target genes (downregulated following *Hnf4a* KO) and **C**) Y-HNF4α-Inhibited target genes (upregulated following *Hnf4a* KO), identified through integration of HNF4α CUT&RUN and hepatocyte-specific *Hnf4a* KO RNA-seq.

**Supplementary Figure 3. Age-associated remodeling of HNF4α target genes disrupts hepatocyte identity and metabolic programs and permits inflammation. A-B**) Gene Ontology Biological Process (GO:BP) enrichment analyses of **A**) Y-HNF4α-activated and **B**) Y-HNF4α-inhibited target genes, stratified by their direction of differential expression in aged wild-type hepatocytes (age-upregulated, age-downregulated, or unchanged). **C-F**) UpSet plots illustrating the overlap of young HNF4α target genes differentially expressed (DE) with age across periportal, midlobular, and pericentral hepatocyte populations, as determined by pseudobulk MERFISH analysis. Genes are stratified by HNF4α regulatory classification and direction of age-associated expression changes: **C**) HNF4α-activated targets downregulated with age, **D**) HNF4α-activated targets upregulated with age, **E**) HNF4α-inhibited targets upregulated with age, and **F**) HNF4α-inhibited targets downregulated with age. Note: genes exhibiting opposing age-associated expression changes across hepatocyte zones are counted in both upregulated and downregulated categories; therefore, summed counts across panels C-F exceed the 122 unique DE HNF4α target genes.

**Supplementary Figure 4. Hepatocyte-specific loss of HNF4α recapitulates and exacerbates key features of liver aging. A**) Median scaled normalized expression of age-associated DE genes identified in WT hepatocytes, represented in whole liver *Hnf4a* KO RNA-seq, stratified by genes upregulated or downregulated with age. **B**) Venn diagram illustrating the overlap of DE genes during physiological aging (O vs Y WT hepatocytes) and following hepatocyte-specific *Hnf4a* deletion in Y mice (Y-KO vs Y-Rosa26 controls). **C-D**) GO:BP enrichment analyses of genes concordantly regulated by physiological aging and following *Hnf4a* deletion in Y, including **C**) genes downregulated in both and **D**) genes upregulated in both. **E**) Median scaled normalized interferon-stimulated gene (ISG) expression in whole liver *Hnf4a* KO RNA-seq. **F**) Heatmap of age-associated expression changes in WT hepatocytes for ISGs identified as direct HNF4α targets. **G**) Representative liver immunohistochemistry images and **H**) quantification for CD45+ (immune cells), F4/80+ (macrophages & Kupffer cells), B220+ (B cells), and CD3+ (T cells). **I-N**) Flow cytometry quantification for the frequency of **I**) CD3+ T cells, **J**) CD4+FoxP3-conventional CD4+ T cells, **K**) CD4+FoxP3+ regulatory T cells, **L**) CD8+ T cells, **M**) F4/80+ macrophages, and **N**) B220+ B cells among CD45hi infiltrating immune cells. Data presented as mean ± [SEM]. n = [number of mice per group]. [one-way ANOVA with Tukey’s test.]

**Supplementary Figure 5. Hepatocyte-specific restoration of HNF4α reverses age-associated liver steatosis and restores FA metabolism gene expression. A)** RT-qPCR results for liver RNA expression of *Hnf4a*. **B**) Whole liver WB and **C**) quantification of HNF4α protein expression in Y and O mice following AAV-mediated HNF4α overexpression or mCherry control. **D**) Liver lipidomics results for total cholesterol. **E**) Volcano plot of differential lipids comparing O-HNF4α versus O-mCherry control. **F**) Log2FC of individual lipid species between O-HNF4α overexpression and O-mCherry control livers, grouped by lipid class. Red dots on E and F indicate significantly altered lipid species (p < 0.05).

