## Supplemental Figures 1-5 & Supplemental Tables 1-2 for "Age-associated repression of HNF4α is a driver and potential therapeutic target in liver metabolic dysfunction"

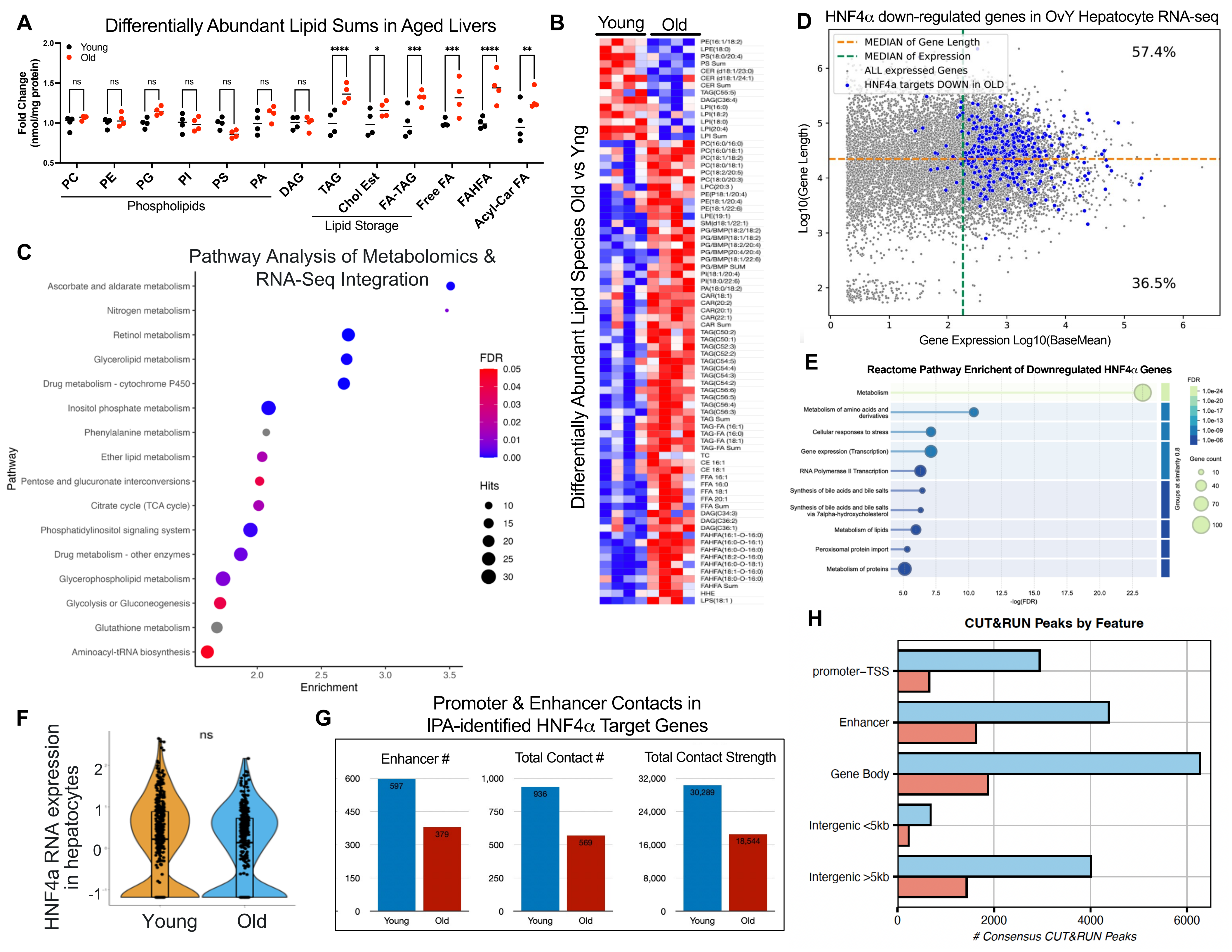
A
Differentially Abundant Lipid Sums in Aged Livers
Young
Old
Fold Change (nmol/mg protein)
ns ns ns ns ns ns ns **** * *** *** **** **
PC PE PG PI PS PA DAG TAG Chol Est FA-TAG Free FA FAHFA Acyl-Car FA
Phospholipids
Lipid Storage
B
Young Old
Differentially Abundant Lipid Species Old vs Yng
C
Pathway Analysis of Metabolomics & RNA-Seq Integration
Ascorbate and aldarate metabolism
Nitrogen metabolism
Retinol metabolism
Glycerolipid metabolism
Drug metabolism - cytochrome P450
Inositol phosphate metabolism
Phenylalanine metabolism
Ether lipid metabolism
Pentose and glucuronate interconversions
Citrate cycle (TCA cycle)
Phosphatidylinositol signaling system
Drug metabolism - other enzymes
Glycerophospholipid metabolism
Glycolysis or Gluconeogenesis
Glutathione metabolism
Aminoacyl-tRNA biosynthesis
Pathway
Enrichment
FDR
Hits
D
HNF4α down-regulated genes in OvY Hepatocyte RNA-seq
MEDIAN of Gene Length
MEDIAN of Expression
ALL expressed Genes
HNF4a targets DOWN in OLD
57.4%
36.5%
Log10(Gene Length)
Gene Expression Log10(BaseMean)
E
Reactome Pathway Enrichent of Downregulated HNF4α Genes
Metabolism
Metabolism of amino acids and derivatives
Cellular responses to stress
Gene expression (Transcription)
RNA Polymerase II Transcription
Synthesis of bile acids and bile salts
Synthesis of bile acids and bile salts via 7alpha-hydroxycholesterol
Metabolism of lipids
Peroxisomal protein import
Metabolism of proteins
-log(FDR)
FDR
Gene count
Groups at similarity 0.8
F
HNF4a RNA expression in hepatocytes
ns
Young Old
G
Promoter & Enhancer Contacts in IPA-identified HNF4α Target Genes
Enhancer #
597 379
Total Contact #
936 569
Total Contact Strength
30,289 18,544
Young Old
H
CUT&RUN Peaks by Feature
promoter–TSS
Enhancer
Gene Body
Intergenic <5kb
Intergenic >5kb
### Consensus CUT&RUN Peaks

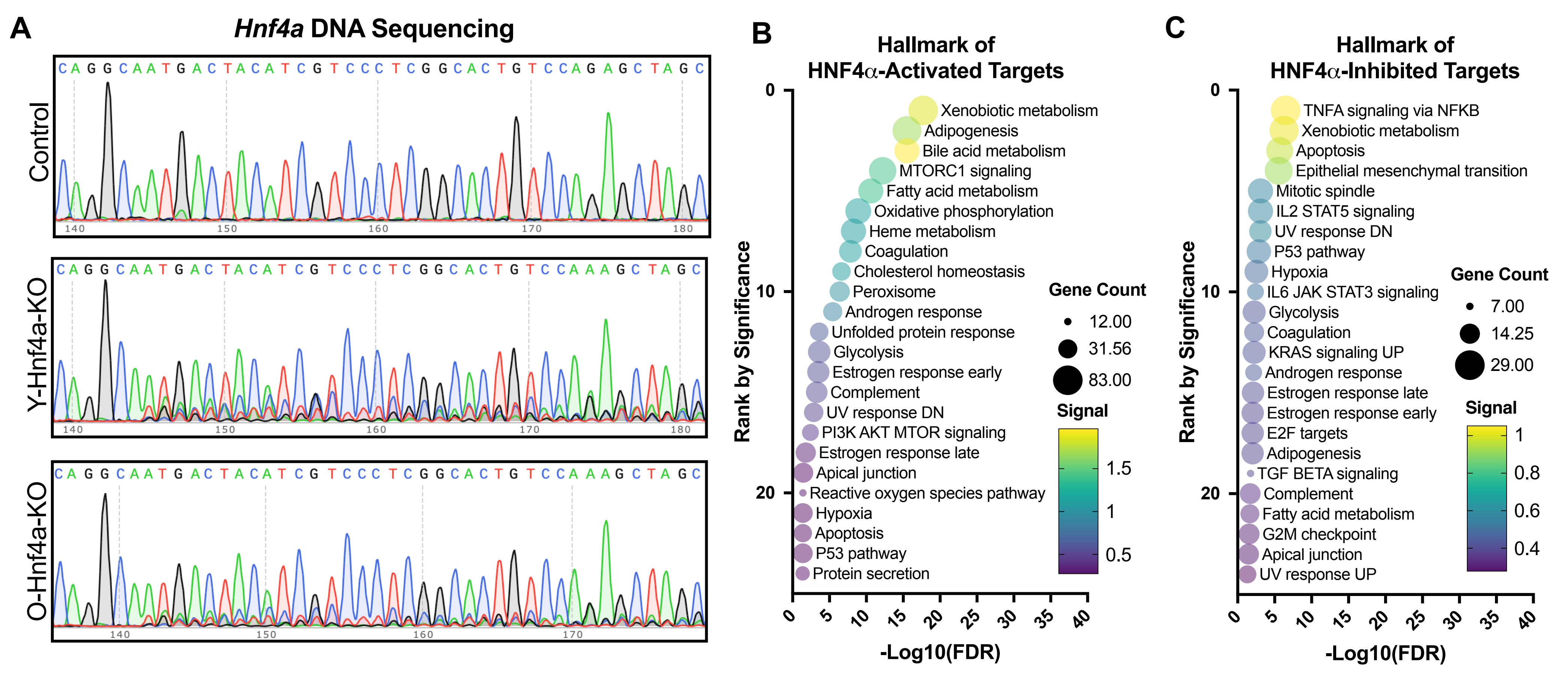

A
Hnf4a DNA Sequencing
Control
CAGGCAATGACTACATCGTCCCTCGGCACTGTCCAGAGCTAGC
140 150 160 170 180
Y-Hnf4a-KO
CAGGCAATGACTACATCGTCCCTCGGCACTGTCCAAAGCTAGC
140 150 160 170 180
O-Hnf4a-KO
CAGGCAATGACTACATCGTCCCTCGGCACTGTCCAAAGCTAGC
140 150 160 170
B
Hallmark of HNF4α-Activated Targets
Rank by Significance
-Log10(FDR)
Xenobiotic metabolism
Adipogenesis
Bile acid metabolism
MTORC1 signaling
Fatty acid metabolism
Oxidative phosphorylation
Heme metabolism
Coagulation
Cholesterol homeostasis
Peroxisome
Androgen response
Unfolded protein response
Glycolysis
Estrogen response early
Complement
UV response DN
PI3K AKT MTOR signaling
Estrogen response late
Apical junction
Reactive oxygen species pathway
Hypoxia
Apoptosis
P53 pathway
Protein secretion
Gene Count
12.00
31.56
83.00
Signal
1.5
1
0.5
C
Hallmark of HNF4α-Inhibited Targets
Rank by Significance
-Log10(FDR)
TNFA signaling via NFKB
Xenobiotic metabolism
Apoptosis
Epithelial mesenchymal transition
Mitotic spindle
IL2 STAT5 signaling
UV response DN
P53 pathway
Hypoxia
IL6 JAK STAT3 signaling
Glycolysis
Coagulation
KRAS signaling UP
Androgen response
Estrogen response late
Estrogen response early
E2F targets
Adipogenesis
TGF BETA signaling
Complement
Fatty acid metabolism
G2M checkpoint
Apical junction
UV response UP
Gene Count
7.00
14.25
29.00
Signal
1
0.8
0.6
0.4

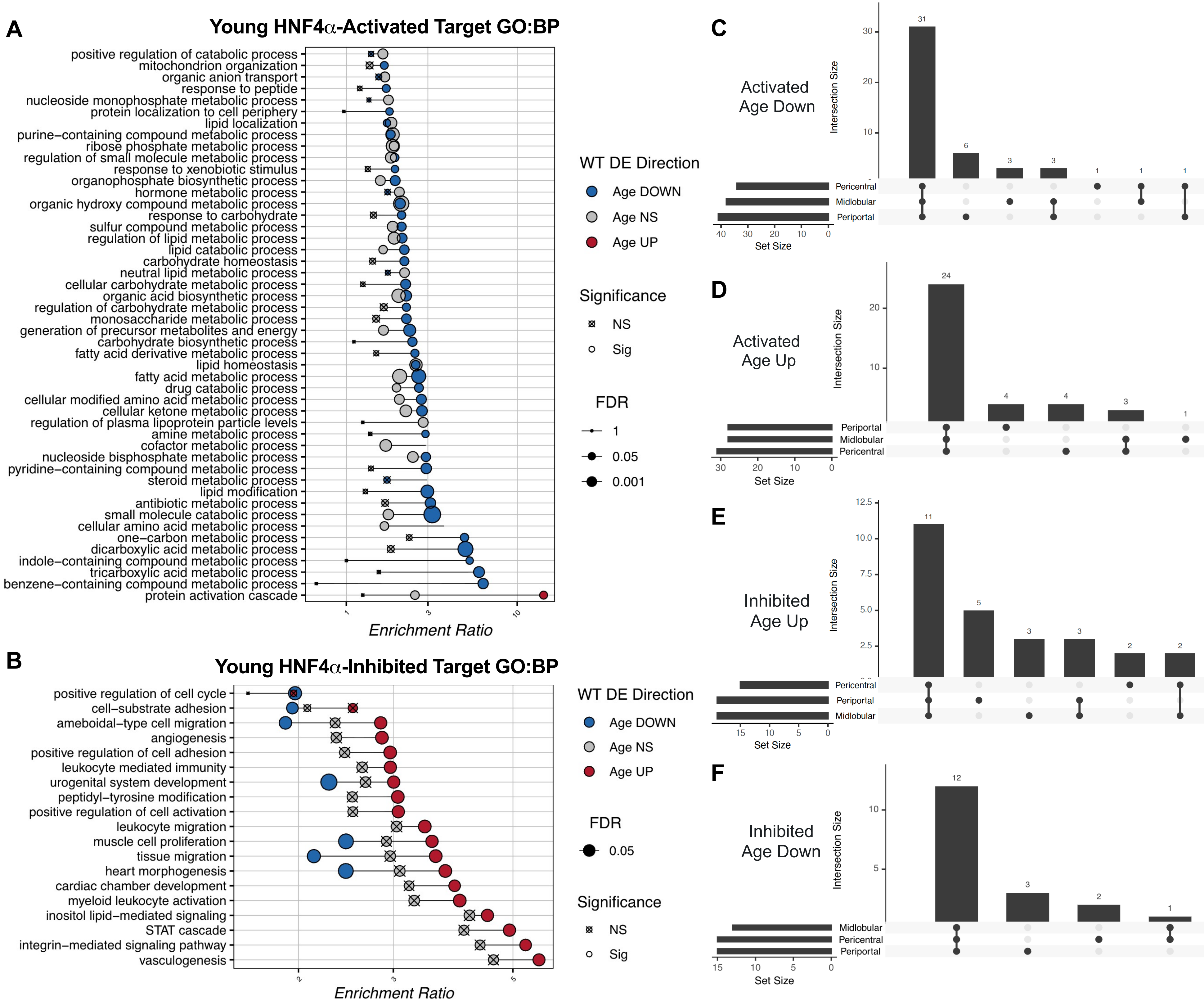
A
Young HNF4α-Activated Target GO:BP
positive regulation of catabolic process
mitochondrion organization
organic anion transport
response to peptide
nucleoside monophosphate metabolic process
protein localization to cell periphery
lipid localization
purine−containing compound metabolic process
ribose phosphate metabolic process
regulation of small molecule metabolic process
response to xenobiotic stimulus
organophosphate biosynthetic process
hormone metabolic process
organic hydroxy compound metabolic process
response to carbohydrate
sulfur compound metabolic process
regulation of lipid metabolic process
lipid catabolic process
carbohydrate homeostasis
neutral lipid metabolic process
cellular carbohydrate metabolic process
organic acid biosynthetic process
regulation of carbohydrate metabolic process
monosaccharide metabolic process
generation of precursor metabolites and energy
carbohydrate biosynthetic process
fatty acid derivative metabolic process
lipid homeostasis
fatty acid metabolic process
drug catabolic process
cellular modified amino acid metabolic process
cellular ketone metabolic process
regulation of plasma lipoprotein particle levels
amine metabolic process
cofactor metabolic process
nucleoside bisphosphate metabolic process
pyridine−containing compound metabolic process
steroid metabolic process
lipid modification
antibiotic metabolic process
small molecule catabolic process
cellular amino acid metabolic process
one−carbon metabolic process
dicarboxylic acid metabolic process
indole−containing compound metabolic process
tricarboxylic acid metabolic process
benzene−containing compound metabolic process
protein activation cascade
1
3
10
Enrichment Ratio
WT DE Direction
Age DOWN
Age NS
Age UP
Significance
NS
Sig
FDR
1
0.05
0.001
B
Young HNF4α-Inhibited Target GO:BP
positive regulation of cell cycle
cell−substrate adhesion
ameboidal−type cell migration
angiogenesis
positive regulation of cell adhesion
leukocyte mediated immunity
urogenital system development
peptidyl−tyrosine modification
positive regulation of cell activation
leukocyte migration
muscle cell proliferation
tissue migration
heart morphogenesis
cardiac chamber development
myeloid leukocyte activation
inositol lipid−mediated signaling
STAT cascade
integrin−mediated signaling pathway
vasculogenesis
2
3
5
Enrichment Ratio
WT DE Direction
Age DOWN
Age NS
Age UP
FDR
0.05
Significance
NS
Sig
C
Activated
Age Down
Intersection Size
30
20
10
31
6
3
3
1
1
1
Pericentral
Midlobular
Periportal
40
30
20
10
0
Set Size
D
Activated
Age Up
Intersection Size
20
10
24
4
4
3
1
Periportal
Midlobular
Pericentral
30
20
10
0
Set Size
E
Inhibited
Age Up
Intersection Size
12.5
10.0
7.5
5.0
2.5
11
5
3
3
2
2
Pericentral
Periportal
Midlobular
15
10
5
0
Set Size
F
Inhibited
Age Down
Intersection Size
10
5
12
3
2
1
Midlobular
Pericentral
Periportal
15
10
5
0
Set Size

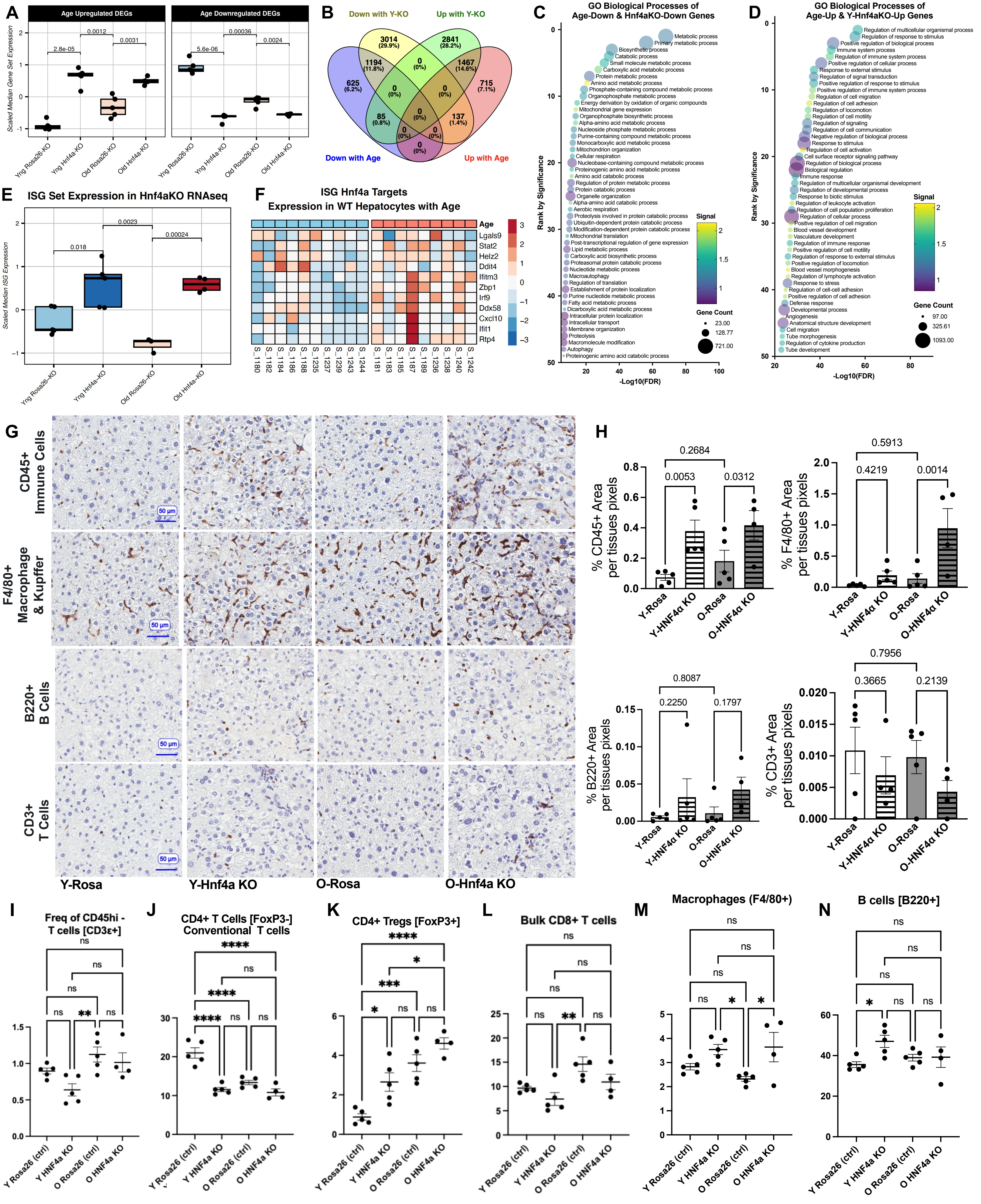

A
Age Upregulated DEGs
Age Downregulated DEGs
Scaled Median Gene Set Expression
2.8e-05
0.0012
0.0031
5.6e-06
0.00036
0.0024
Yng Rosa26-KO
Yng Hnf4a-KO
Old Rosa26-KO
Old Hnf4a-KO
B
Down with Y-KO
Up with Y-KO
Down with Age
Up with Age
3014 (29.9%)
2841 (28.2%)
1194 (11.8%)
1467 (14.6%)
625 (6.2%)
715 (7.1%)
85 (0.8%)
137 (1.4%)
0 (0%)
C
GO Biological Processes of
Age-Down & Hnf4aKO-Down Genes
Rank by Significance
-Log10(FDR)
Signal
Gene Count
23.00
128.77
721.00
Metabolic process
Primary metabolic process
Biosynthetic process
Catabolic process
Small molecule metabolic process
Carboxylic acid metabolic process
Protein metabolic process
Amino acid metabolic process
Phosphate-containing compound metabolic process
Organophosphate metabolic process
Energy derivation by oxidation of organic compounds
Mitochondrial gene expression
Organophosphate biosynthetic process
Alpha-amino acid metabolic process
Nucleoside phosphate metabolic process
Purine-containing compound metabolic process
Monocarboxylic acid metabolic process
Mitochondrion organization
Cellular respiration
Nucleobase-containing compound metabolic process
Proteinogenic amino acid metabolic process
Amino acid catabolic process
Regulation of protein metabolic process
Protein catabolic process
Organelle organization
Alpha-amino acid catabolic process
Aerobic respiration
Proteolysis involved in protein catabolic process
Ubiquitin-dependent protein catabolic process
Modification-dependent protein catabolic process
Mitochondrial translation
Post-transcriptional regulation of gene expression
Lipid metabolic process
Carboxylic acid biosynthetic process
Proteasomal protein catabolic process
Nucleotide metabolic process
Macroautophagy
Regulation of translation
Establishment of protein localization
Purine nucleotide metabolic process
Fatty acid metabolic process
Dicarboxylic acid metabolic process
Intracellular protein localization
Intracellular transport
Membrane organization
Proteolysis
Macromolecule modification
Autophagy
Proteinogenic amino acid catabolic process
D
GO Biological Processes of
Age-Up & Y-Hnf4aKO-Up Genes
Rank by Significance
-Log10(FDR)
Signal
Gene Count
97.00
325.61
1093.00
Regulation of multicellular organismal process
Regulation of response to stimulus
Positive regulation of biological process
Immune system process
Regulation of immune system process
Positive regulation of cellular process
Response to external stimulus
Regulation of signal transduction
Positive regulation of response to stimulus
Positive regulation of immune system process
Regulation of cell migration
Regulation of cell adhesion
Regulation of locomotion
Regulation of cell motility
Regulation of signaling
Regulation of cell communication
Negative regulation of biological process
Response to stimulus
Regulation of cell activation
Cell surface receptor signaling pathway
Regulation of biological process
Biological regulation
Immune response
Regulation of multicellular organismal development
Regulation of developmental process
Response to biotic stimulus
Regulation of leukocyte activation
Regulation of cell population proliferation
Regulation of cellular process
Positive regulation of cell migration
Blood vessel development
Vasculature development
Regulation of immune response
Positive regulation of cell motility
Regulation of response to external stimulus
Positive regulation of locomotion
Blood vessel morphogenesis
Regulation of lymphocyte activation
Response to stress
Regulation of cell-cell adhesion
Positive regulation of cell adhesion
Defense response
Developmental process
Angiogenesis
Anatomical structure development
Cell migration
Tube morphogenesis
Regulation of cytokine production
Tube development
E
ISG Set Expression in Hnf4aKO RNAseq
Scaled Median ISG Expression
0.018
0.0023
0.00024
Yng Rosa26-KO
Yng Hnf4a-KO
Old Rosa26-KO
Old Hnf4a-KO
F
ISG Hnf4a Targets
Expression in WT Hepatocytes with Age
Age
Lgals9
Stat2
Helz2
Ddit4
Ifitm3
Zbp1
Irf9
Ddx58
Cxcl10
Ifit1
Rtp4
S_180 S_182 S_184 S_186 S_188 S_235 S_237 S_239 S_243 S_244 S_181 S_183 S_185 S_187 S_189 S_236 S_238 S_240 S_242
G
CD45+ Immune Cells
F4/80+ Macrophage & Kupffer
B220+ B Cells
CD3+ T Cells
50 µm
Y-Rosa
Y-Hnf4a KO
O-Rosa
O-Hnf4a KO
H
% CD45+ Area per tissues pixels
0.2684
0.0053
0.0312
% F4/80+ Area per tissues pixels
0.5913
0.4219
0.0014
% B220+ Area per tissues pixels
0.8087
0.2250
0.1797
% CD3+ Area per tissues pixels
0.7956
0.3665
0.2139
Y-Rosa
Y-HNF4α KO
O-Rosa
O-HNF4α KO
I
Freq of CD45hi - T cells [CD3ε+]
J
CD4+ T Cells [FoxP3-] Conventional T cells
K
CD4+ Tregs [FoxP3+]
L
Bulk CD8+ T cells
M
Macrophages (F4/80+)
N
B cells [B220+]
ns
*
**
***
****
Y Rosa26 (ctrl)
Y HNF4a KO
O Rosa26 (ctrl)
O HNF4a KO

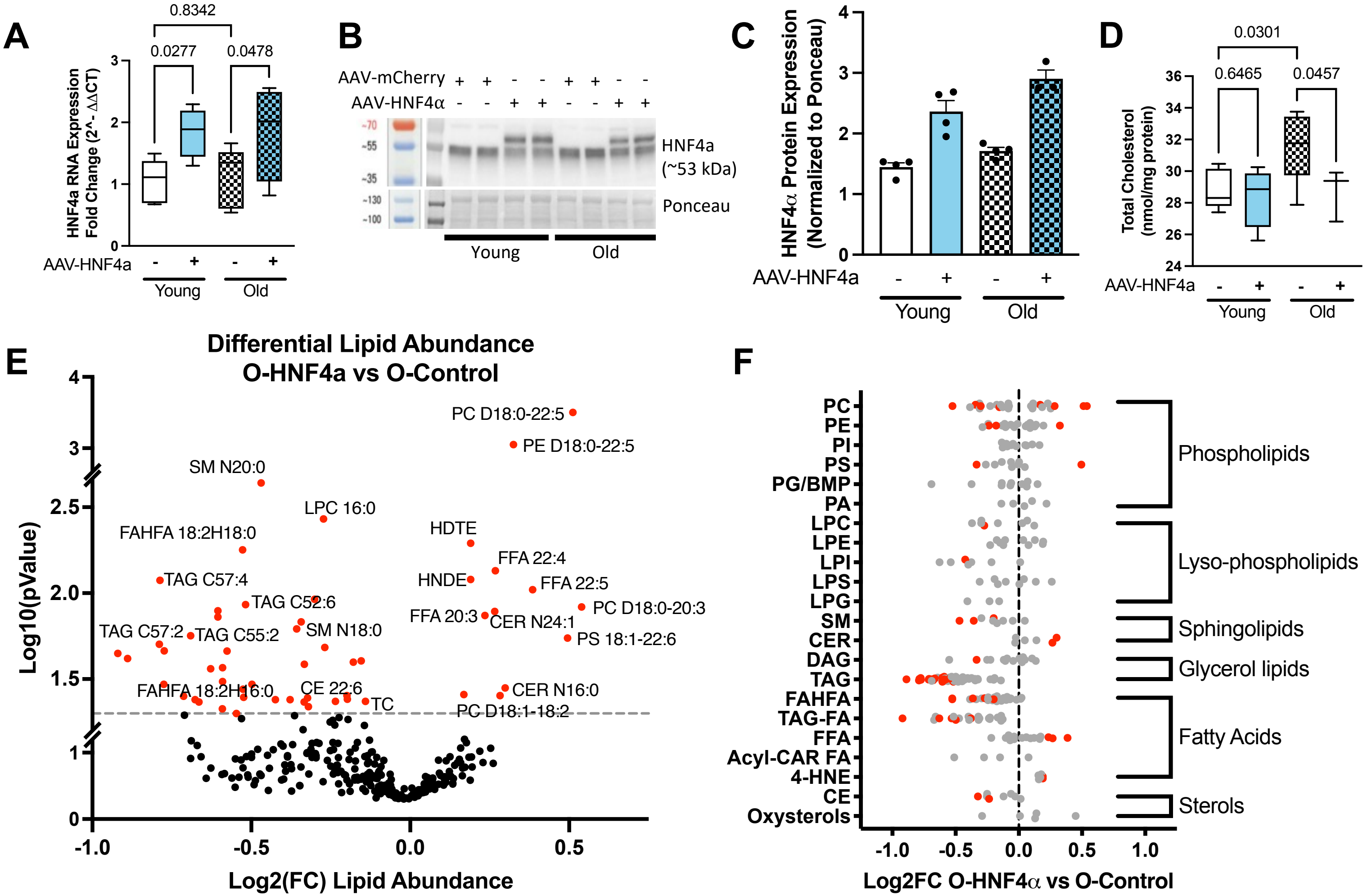
A
HNF4a RNA Expression Fold Change (2^- ΔΔCT)
0.8342
0.0277
0.0478
AAV-HNF4a
Young
Old
B
AAV-mCherry
AAV-HNF4α
HNF4a (~53 kDa)
Ponceau
Young
Old
C
HNF4α Protein Expression (Normalized to Ponceau)
AAV-HNF4a
Young
Old
D
Total Cholesterol (nmol/mg protein)
0.0301
0.6465
0.0457
AAV-HNF4a
Young
Old
E
Differential Lipid Abundance
O-HNF4a vs O-Control
Log10(pValue)
Log2(FC) Lipid Abundance
PC D18:0-22:5
PE D18:0-22:5
SM N20:0
LPC 16:0
FAHFA 18:2H18:0
HDTE
FFA 22:4
TAG C57:4
HNDE
FFA 22:5
TAG C52:6
PC D18:0-20:3
FFA 20:3
CER N24:1
SM N18:0
TAG C57:2
TAG C55:2
PS 18:1-22:6
FAHFA 18:2H16:0
CE 22:6
TC
CER N16:0
PC D18:1-18:2
F
PC
PE
PI
PS
PG/BMP
PA
LPC
LPE
LPI
LPS
LPG
SM
CER
DAG
TAG
FAHFA
TAG-FA
FFA
Acyl-CAR FA
4-HNE
CE
Oxysterols
Phospholipids
Lyso-phospholipids
Sphingolipids
Glycerol lipids
Fatty Acids
Sterols
Log2FC O-HNF4α vs O-Control

**Supplementary Tables 1: Antibodies used for Flow Cytometry - *T cell panel***

| Marker | Clone | Conjugate | Vendor | Catalog no. | Dilution |
|---|---|---|---|---|---|
| **CD8α** | 53-6.7 | BUV395 | BD | 563786 | 1/200 |
| **CD44** | IM7 | BUV496 | BD | 741057 | 1/200 |
| **CD4** | RM4-5 | BUV737 | BD | 612843 | 1/200 |
| **IFNγ** | XMG1.2 | BUV805 | BD | 569625 | 1/100 |
| **CX3CR1** | SA011F11 | BV421 | BioLegend | 149023 | 1/200 |
| **SLAMF6** | 13G3 | BV605 | BD | 745250 | 1/200 |
| **CD45** | 30-F11 | BV650 | Invitrogen | 416-0451-82 | 1/200 |
| **TIM3** | RMT3-23 | BV711 | BioLegend | 119727 | 1/200 |
| **PD-1** | 29F.1A12 | BV785 | BioLegend | 135225 | 1/200 |
| **TCF1** | C63D9 | AF488 | Cell Signaling Technology | 6444S | 1/100 |
| **TNFα** | MP6-XT22 | PerCP/eF710 | Invitrogen | 46-7321-82 | 1/100 |
| **TOX** | TXRX10 | PE | Invitrogen | 12-6502-82 | 1/100 |
| **LIVE/DEAD** | — | Fixable Red | Invitrogen | L34972 | 1/1,000 |
| **CD101** | Moushi101 | PE/Cy7 | Invitrogen | 25-1011-82 | 1/200 |
| **FoxP3** | FJK-16s | APC | Invitrogen | 17-5773-82 | 1/100 |
| **Ki67** | SolA15 | AF700 | Invitrogen | 56-5698-82 | 1/100 |
| **Granzyme B** | QA16A02 | APC/Fire 750 | BioLegend | 372210 | 1/100 |

**Supplementary Table 2: Antibodies used for Flow Cytometry - *Myeloid panel***

| Marker | Clone | Conjugate | Vendor | Catalog no. | Dilution |
|---|---|---|---|---|---|
| **Ly6G** | 1A8 | BUV395 | BD | 563978 | 1/200 |
| **CD11b** | M1/70 | BUV737 | BD | 612800 | 1/500 |
| **MHC-II** | M5/114.15.2 | BV421 | BioLegend | 107632 | 1/500 |
| **Ly6C** | HK1.4 | BV605 | BioLegend | 128036 | 1/500 |
| **CD45** | 30-F11 | BV650 | Invitrogen | 416-0451-82 | 1/200 |
| **CD3ε** | 17A2 | BV711 | BioLegend | 100241 | 1/100 |
| **CD11c** | N418 | BV785 | BioLegend | 117336 | 1/100 |
| **BST2** | 927 | FITC | BioLegend | 127008 | 1/200 |
| **NK1.1** | PK136 | PerCP/Cy5.5 | BioLegend | 108728 | 1/200 |
| **PD-L1** | 10F.9G2 | PE | BioLegend | 124308 | 1/200 |
| **LIVE/DEAD** | — | Fixable Red | Invitrogen | L34972 | 1/1,000 |
| **CX3CR1** | SA011F11 | PE/Cy7 | BioLegend | 149016 | 1/200 |
| **CD206** | C068C2 | APC | BioLegend | 141710 | 1/200 |
| **B220** | RA3-6B2 | AF700 | BioLegend | 103232 | 1/200 |
| **F4/80** | BM8 | APC/Cy7 | BioLegend | 123116 | 1/100 |
